# *SALL3* mediates the loss of neuroectodermal differentiation potential in human embryonic stem cells with chromosome 18q loss

**DOI:** 10.1101/2023.06.26.546513

**Authors:** Yingnan Lei, Diana Al Delbany, Nuša Krivec, Marius Regin, Edouard Couvreu de Deckersberg, Charlotte Janssens, Manjusha Ghosh, Karen Sermon, Claudia Spits

## Abstract

Human pluripotent stem cell (hPSC) cultures are prone to genetic drift, as cells that have acquired specific genetic abnormalities experience a selective advantage in vitro. These abnormalities are highly recurrent in hPSC lines worldwide, but currently their functional consequences in differentiating cells are scarcely described. An accurate assessment of the risk associated with these genetic variants in both research and clinical settings is therefore lacking. In this work, we established that one of these recurrent abnormalities, the loss of chromosome 18q, impairs neuroectoderm commitment and affects the cardiac progenitor differentiation of hESCs. We show that downregulation of *SALL3*, a gene located in the common 18q loss region, is responsible for failed neuroectodermal differentiation. Knockdown of *SALL3* in control lines impaired differentiation in a manner similar to the loss of 18q, while transgenic overexpression of *SALL3* in hESCs with 18q loss rescued the differentiation capacity of the cells. Finally, we show by gene expression analysis that loss of 18q and downregulation of *SALL3* leads to changes in the expression of genes involved in pathways regulating pluripotency and differentiation, including the WNT, NOTCH, JAK-STAT, TGF-beta and NF-kB pathways, suggesting that these cells are in an altered state of pluripotency.

## Introduction

Human pluripotent stem cells (hPSCs), including human embryonic stem cells (hESCs) and induced pluripotent stem cells (hiPSCs), are self-renewing cells that can give rise to any cell type originating from any of the three embryonic germ layers^1, 2^. This makes hPSCs an attractive resource for *in vitro* disease modelling, developmental biology research, drug discovery, and cell transplantation therapy. A substantial number of clinical trials are underway using hPSC-derived cell products, including for the treatment of age-related macular degeneration, spinal cord injury and type 1 diabetes^3–5^. An important hurdle for both safe clinical translation and the reliable use of hPSCs as *in vitro* research models is the occurrence of cell culture drift due to the acquisition of genetic abnormalities^5–8^. A subset of these genetic aberrations are highly recurrent and are found in hPSC lines worldwide. These recurrent changes vary in size from single-nucleotide point mutations to large chromosome structural variants, the most common being gains of chromosomes 1q, 12p, 17, 20, and X and losses of 10p, 18q and 22p, as well as mutations in TP53^6, 7, 9–12^. Other genetic changes include epigenetic variations, including erosion of X-chromosome inactivation^13, 14^, mutations in the mitochondrial genome^15, 16^ and an array of other point mutations and structural variants spread throughout the genome^10, 11^

These recurrent genetic changes arise from the pool of common variations in hPSC cultures via different cell competition mechanisms. hPSCs are prone to replication stress, leading to DNA damage, which in turn is a source of de novo genetic variation^17–21^. For instance, up to 20% of cells in an hESC culture carry de novo structural variants, but only a minority of them have the potential to confer a selective advantage to the cells^20, 22, 23^, leading to the mutant cells rapidly outcompeting their genetically balanced counterparts^24–27^. The exact traits that the different chromosomal abnormalities confer on undifferentiated cells, as well as the specific driver genes of these traits, are only well established for the gain of 20q11.21^28^. This abnormality confers a decreased sensitivity to apoptosis-inducing events due to increased expression of the gene *BCL2L1*, located in the minimal gained region^6, 26, 27^. For gains of 12p, it is considered that *NANOG* drives at least part of the growth advantage of the cells^29^, and cells with a complex karyotype carrying all the most common abnormalities (gains of 1, 12, 17 and 20q) can outcompete the other cells by corralling and mechanical compression^24^.

An important concern about these genetic variants is whether and how they alter the differentiation capacity of hPSCs and potentially prime differentiated cells for malignant transformation^8, 30–32^. The gain of 1q is common in cancers, particularly lung adenocarcinoma, breast invasive carcinoma and liver hepatocellular carcinoma^33^, and appears to confer a growth advantage to cells during differentiation from hESCs to neural precursors^34^. Moreover, variants in 1q21.1 can alter neurodevelopmental trajectories upon hiPSC differentiation, with the deletion of 1q21.1 accelerating neuronal production and its duplication delaying the transition from neural progenitor cell to neuron^35^. Gains in chromosome 12 are frequently found in testicular germ cell tumors^36^. hPSCs with trisomy 12 display a reduced tendency toward spontaneous differentiation^29, 37^, and gain of 12p13.31 results in an overall reduction in trilineage differentiation capacity and foci of residual pluripotent cells during hepatic differentiation^38, 39^. The highly recurrent gain of 20q11.21 impairs the neuroectodermal lineage commitment of hPSCs^40, 41^, and hESCs with 20q11.1q11.2 amplification have a reduced propensity to differentiate down the hematopoietic lineage, maintain more immature phenotypes along the neural differentiation trajectory and generate teratomas with foci of undifferentiated cells^42^. The gain of chromosome 17 is common in neuroblastomas, testicular germ cell tumors and breast cancers^36^. hPSC lines with a gain of chromosome 17 show altered differentiation patterns in embryoid bodies^43^, and the amplification of *WNT3A* and *WNT9* on chromosome 17q21.31 alters neuronal differentiation^44^.

Deletions of chromosome 18q are one of the rarer recurrent structural chromosomal abnormalities in hPSCs. This deletion was first reported as a single event by Maitra *et al.* in 2005^45^, and our group later found 18q deletions in three different hESC lines at relatively early passages, always as part of a derivative chromosome 18^28^. A large study by Amps *et al*. in 2011 revealed 5 instances of this deletion^46^, A large study by Amps *et al*. in 2011 revealed 5 instances of this deletion, and WiCell has reported that 4% of the 7300 hPSC cultures evaluated over nearly eight years carried an 18q deletion (WiCell Cytogenetics Lab, https://www.wicell.org/media.acux/29102c0e-e88e-426b-ab7d-bac4c2a9ec6a). Chromosome 18q loss is common in cancers, especially gastrointestinal tract cancers^33^, and is linked to several disorders, including congenital malformations, developmental delays, and intellectual disability^47^. However, the impact of 18q loss on the functional properties of hPSCs is unknown. Therefore, the aim of this work was to examine the functional effects of 18q deletions during hPSC differentiation into the three embryonic germ layers and to determine the molecular mechanisms involved.

## RESULTS

### The minimal common region of 18q loss spans 14 genes expressed in undifferentiated hESCs and includes *SALL3*

We initially identified 18q losses in the hESC lines VUB04 and VUB26, in the form of a derivative chromosome 18. VUB04 presented a deletion at 18q21.2qter and a duplication at 5q14.2qter, and VUB26 showed the minimal 18q loss region (18q23qter) and a duplication at 7q33qter (Fig 1A. Sup. Fig. 1 and Sup. Table 1)^48^. These two hESC lines were not used in the present work because they further genetically drifted and acquired gain of 1q and 20q11.21, but their analysis helped to narrow down the common 18q deletion region. In this study, we used two other hESC lines bearing derivative chromosomes 18 involving a loss of 18q losses (hESC^del18q^), VUB14^del18q^ and VUB13^del18q^, as well as three chromosomally balanced lines (hESC^WT^), VUB14^WT^, and VUB04^WT^ and VUB03^WT^, which served as controls for VUB13^del18q^ since VUB13^WT^ was lost (details on the karyotypes of the lines and their characterization are shown in Sup. Fig. 1 and Sup. Table 1). We used shallow genome sequencing to confirm the karyotypes before starting the experiments and all lines were routinely inspected with qPCR assays targeting recurrent chromosomal abnormalities (1q, 12p, 20q11.21 and 17q) to confirm their genomic stability for the duration of the different experiments.

**Figure 1.**
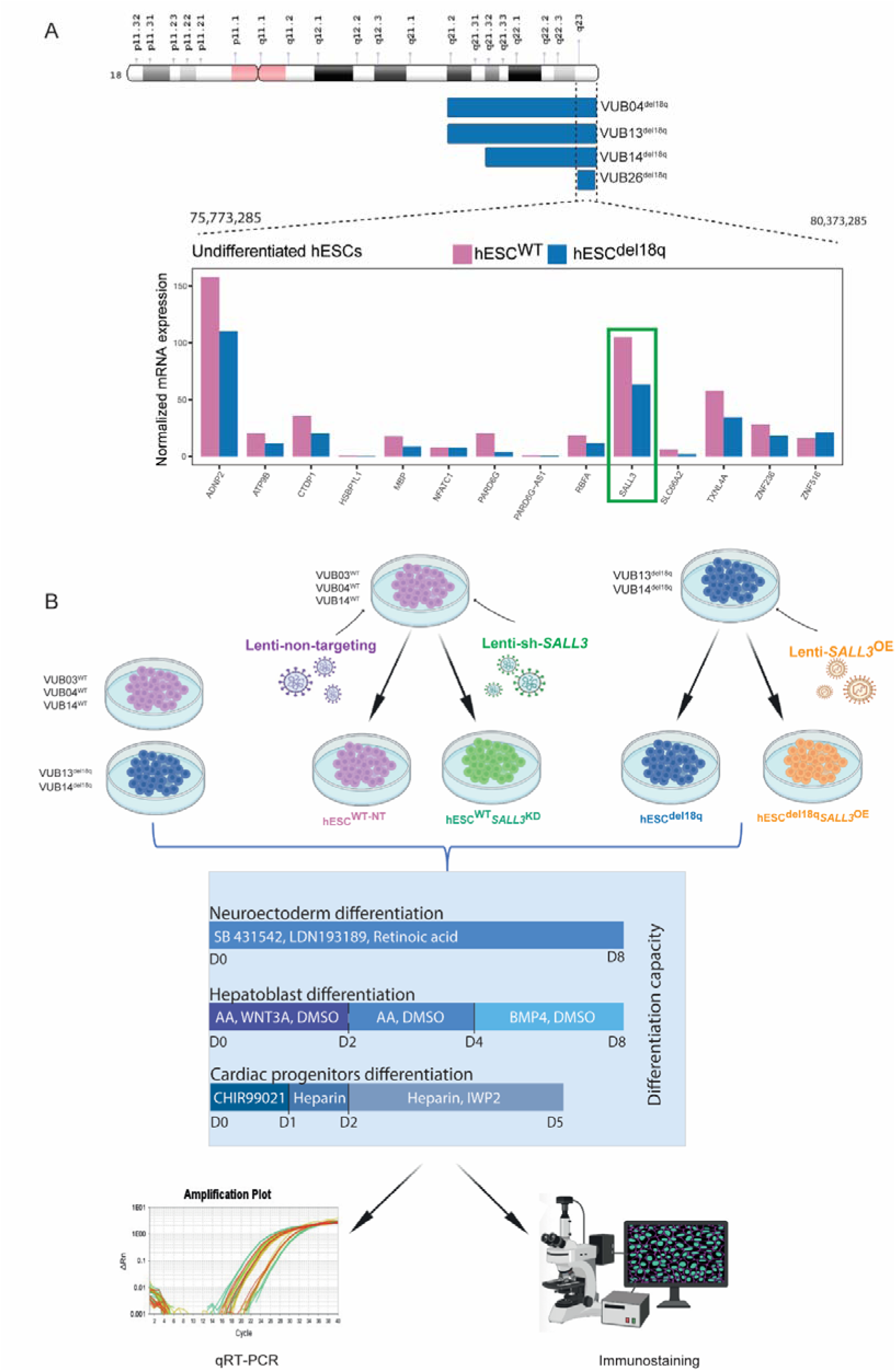
The minimal common region of 18q loss spans 14 genes expressed in undifferentiated hESCs, including *SALL3*. (A) Graphical representation of the regions of chromosome 18 loss in all VUB hESC^del18q^ lines and showing the normalized expression of the genes located in the minimal common region of loss examined by bulk-RNA sequencing to compare the transcriptome of hESC^del18q^ and hESC^WT^ as well as the genetically modified counterparts. (B) Overview of the experimental setup. We used two lines carrying a loss of 18q and three genetically balanced lines. The WT lines were also genetically modified to down-regulate *SALL3*, while 18q loss lines were modified to over-express *SALL3*. All lines were subjected to differentiate to neuroectoderm, hepatoblasts and cardiac progenitors, all differentiation experiments were carried out at least in triplicate. qRT-PCR and immunostaining were used to assess the identity of the differentiated cells. AA: Activin A; Lenti-sh*-SALL3*: lentiviral transduction for a short-hairpin RNA against *SALL3.* Lenti-O3-*SALL3*: lentiviral transduction with a construct for the transgenic expression of *SALL3*.

The 18q losses exhibited a common loss region from bp 75,773,285 to 80,373,285, spanning 37 loci. Bulk RNA sequencing of undifferentiated hESCs indicated that 14 genes within this region are expressed in undifferentiated hESCs, with counts per million greater than one in at least two samples (Fig. 1A). Of these coding genes, *ADNP2*, *SALL3* and *TXNL4A* had the highest expression and showed decreased transcript levels in mutant cells. *ADNP2* and *TXNL4A* have no known function in hPSC. *ADNP2* is predicted to be a transcription factor, and its silencing increases oxidative stress-mediated cell death^49^. *TXNL4A* is a component of the U5 small ribonucleoprotein particle, which is involved in pre-mRNA splicing and is associated with Burn-McKeown syndrome^50^. *SALL3* was more promising as candidate driver gene as it has previously been reported to regulate the differentiation propensity of hiPSC lines^51^. Kuroda *et al*. showed that hiPSC lines expressing high levels of *SALL3* differentiated preferentially into ectoderm, while hiPSC lines expressing lower levels of *SALL3* tended to differentiate into mesoderm and endoderm^51^. It has also been shown that *SALL3* interacts with the Mediator complex in neural stem cells^52^ and is related to the development of the nervous system^53^. Considering these previous findings, we hypothesized that decreased expression of *SALL3* as a result of a loss of one copy of the gene could alter the differentiation capacity of hESCs^del18q^.

### hESCs with 18q loss show impaired neuroectoderm differentiation

As a first step, we investigated the effect of 18q deletion on hESC ectoderm lineage commitment. hESC^WT^ and hESC^del18q^ were subjected to neuroectoderm differentiation for 8 days using LDN193189 (LDN), SB431542 (SB) and retinoic acid (RA)^54^ (Fig. 1B). We measured the mRNA levels of different neuroectoderm markers to evaluate neuroectoderm differentiation efficiency and the expression of the undifferentiated state markers *NANOG* and *POUF51* (Fig. 2A and Sup Fig. 2A). VUB13^del18q^ and VUB14^del18q^ had significantly lower mRNA levels of *PAX6*, *NES* and *SOX1*, which were decreased by 3-fold, 15-fold, and 25-fold, respectively, compared to the levels in hESC^WT^ (p≤0.0001 unpaired t-test), indicating a decreased neuroectodermal differentiation efficiency in hESC^del18q^. *POU5F1* and *NANOG* mRNA expression levels were almost undetectable for all differentiated cells (Sup Fig. 2A). We also evaluated the differentiation of hESC^WT^ and hESC^del18q^ cells by immunostaining (Fig. 2B-C). We observed a lower percentage of PAX6 positive cells in differentiated hESC^del18q^ than in differentiated hESC^WT^ cells (45% vs 70%, p = 0.0114, unpaired t-test, Fig. 2C), which was consistent with the decrease in the levels of *PAX6* mRNA. Taken together, these results show that hESCs^del18q^ differentiation into neuroectoderm is impaired, and rather than to remain undifferentiated state, they miss-specify.

**Figure 2.**
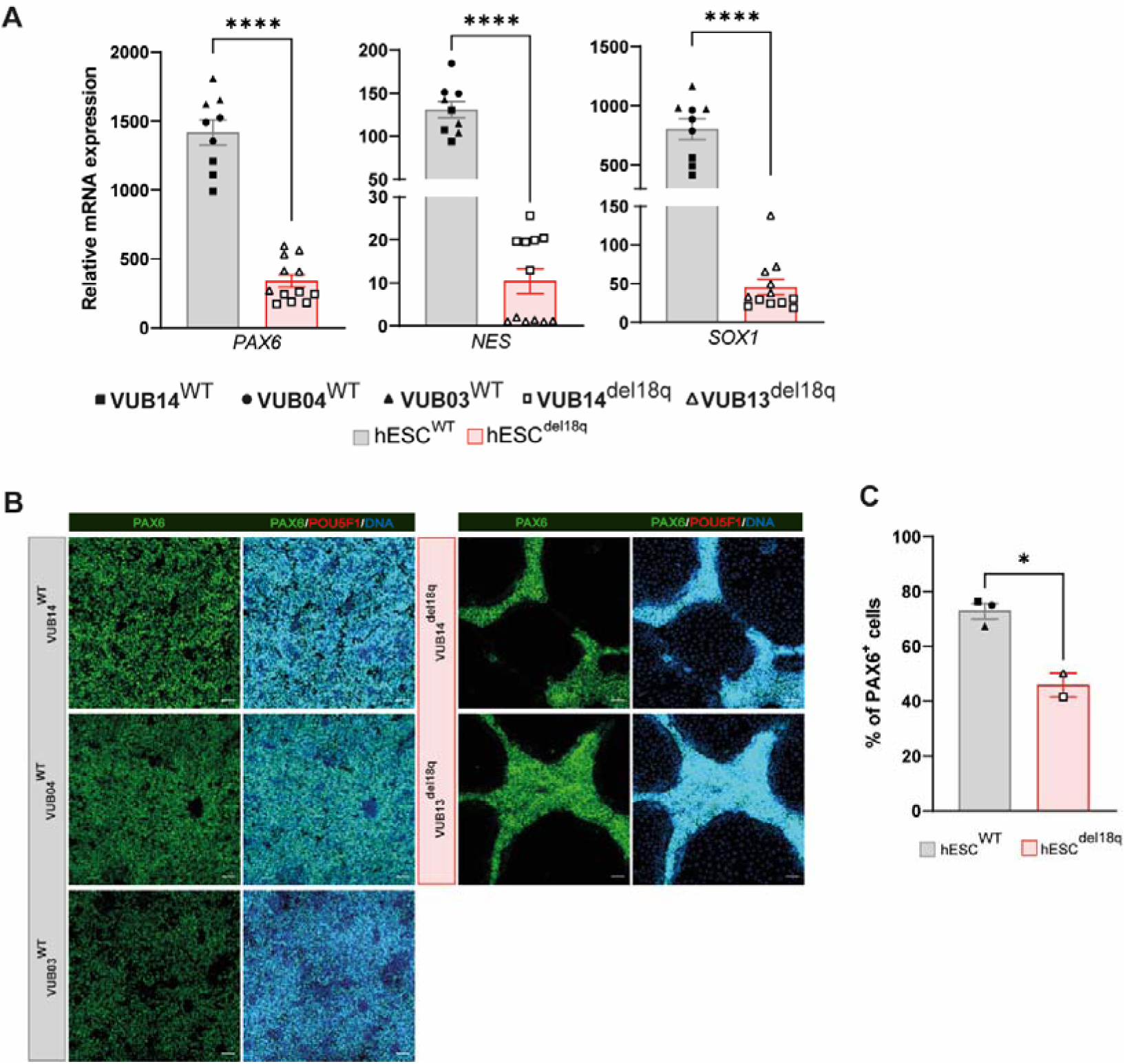
hESCs with 18q loss show impaired neuroectoderm differentiation. (A) Relative mRNA expression as measured by qRT-PCR for ectoderm markers *PAX6*, *NES* and *SOX1* (n = 3–6). Data are shown as the means ± SEM. Different patterns indicate different cell lines, and the horizontal bars with asterisks *, **, *** and **** represent statistical significance between samples at 5%, 1%, 0.1% and 0.01% respectively (unpaired t-test). (B) Immunostaining for PAX6 (green) and POU5F1 (red) in mutant and control lines after 8 days of neuroectoderm differentiation, all scale bars are 50 µm. (C) Percentage of PAX6-positive cells in the immunostainings shown in B.

### hESC^del18q^ readily differentiates into mesendoderm derivatives but shows abnormal cardiomyocyte progenitor differentiation

We further investigated the impact of 18q deletion on the mesendoderm differentiation capacity of hESCs by differentiating hESC^WT^ and hESC^del18q^ into, in this case cardiac progenitors and hepatoblasts. Thus, we induced the differentiation of hESC^WT^ and hESC^del18q^ into mesoderm using a 5-day cardiac progenitor induction protocol as described previously^55^ (Fig. 1B). We evaluated the mRNA levels of the cardiac progenitor markers *GATA4*, *ISL1*, *NKX2-5* and *PDGFRA* (Fig. 3A). hESC^del18q^ showed 2-fold lower levels of *GATA4* mRNA (p = 0.0012 unpaired t-test) (Fig. 3A). hESC^del18q^ lines expressed *ISL1* at 15-fold higher levels, on average (p=0.0016, unpaired t-test), than hESC^WT^ lines. We found no significant difference in the expression levels of *NKX2-5* or *PDGFRA* between the control and mutant cardiac progenitor groups (p=0.11 and p=0.42, unpaired t-test, Fig. 3A). We also evaluated the proportion of differentiated cardiac progenitor and undifferentiated cells by immunostaining for GATA4 and POU5F1. The percentage of GATA4-positive cells was not statistically significantly different in hESC^del18q^ than in hESC^WT^ (p=0.2942, unpaired t-test, Fig. 3B-C), mainly due to the low differentiation efficiency of VUB03^WT^, while the cells were all POU5F1 negative. Taken together, and bearing in mind the temporal expression of these markers during cardiac differentiation^56, 57^, the results suggest that hESC^del18q^ may experience differentiation delays or arrest, reaching an ISL1^high^ GATA4^low^ stage at day 5 but remaining less mature than their hESC^WT^ counterparts.

**Figure 3.**
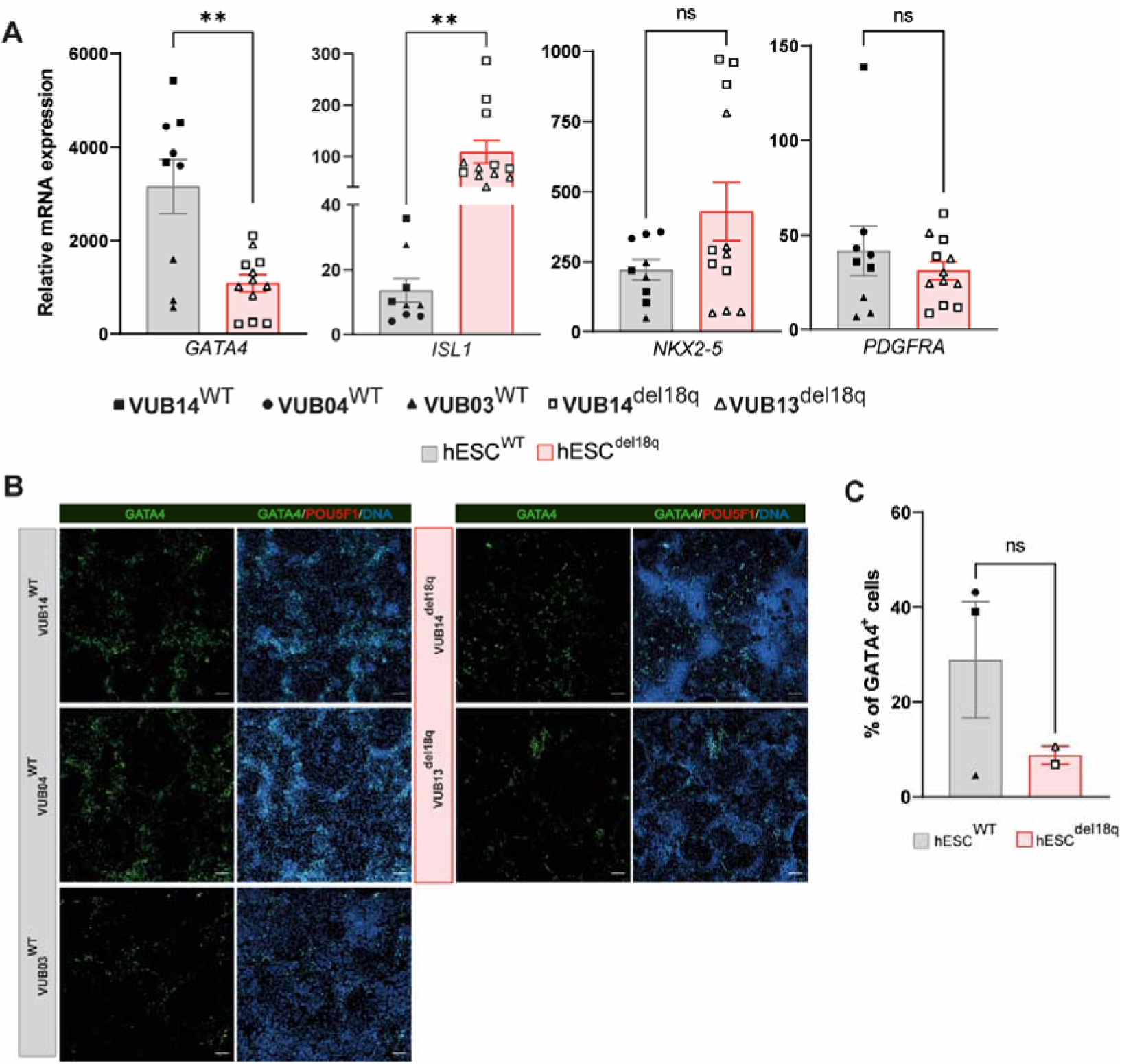
hESC^del18q^ cells show abnormal cardiomyocyte progenitor differentiation. (A) Relative mRNA expression as measured by qRT-PCR for the cardiac progenitor markers *GATA4*, *ISL1*, *NKX2-5* and *PDGFRA* (n = 3–6). Data are shown as the means ± SEM; different patterns indicate different cell lines, and the horizontal bars with asterisks *, **, *** and **** represent statistical significance between samples at 5%, 1%, 0.1% and 0.01% respectively (unpaired t-test). (B) Immunostaining for NKX2-5 (green) and POU5F1 (red) in mutant and control lines after 5 days of cardiac progenitor differentiation, all scale bars are 100 µm. (C) Percentage of GATA4-positive cells in the immunostainings shown in B.

Next, we differentiated hESC^WT^ and hESC^del18q^ into hepatoblasts by applying the modified differentiation protocol for 8 days as described previously^58^ (Fig. 1B). We measured the expression levels of the hepatoblast markers *HNF4A*, *AFP*, *ALB* and *FOXA2* (Fig. 4A). We found no significant difference in the mRNA expression levels of *HNF4A*, *ALB*, and *FOXA2* between hESC^WT^ and hESC^del18q^ (p>0.05, unpaired t-test), while AFP had a lower expression in hESC^del18q^ (Fig. 4A, p=0.0095, unpaired t-test). We further evaluated hepatoblast differentiation by immunostaining for HNF4A and found that the percentages of HNF4A^-^positive cells in hESC^del18q^ cells were similar to those in the wild type (Fig. 4B-C, p=0.2514, unpaired t-test).

**Figure 4.**
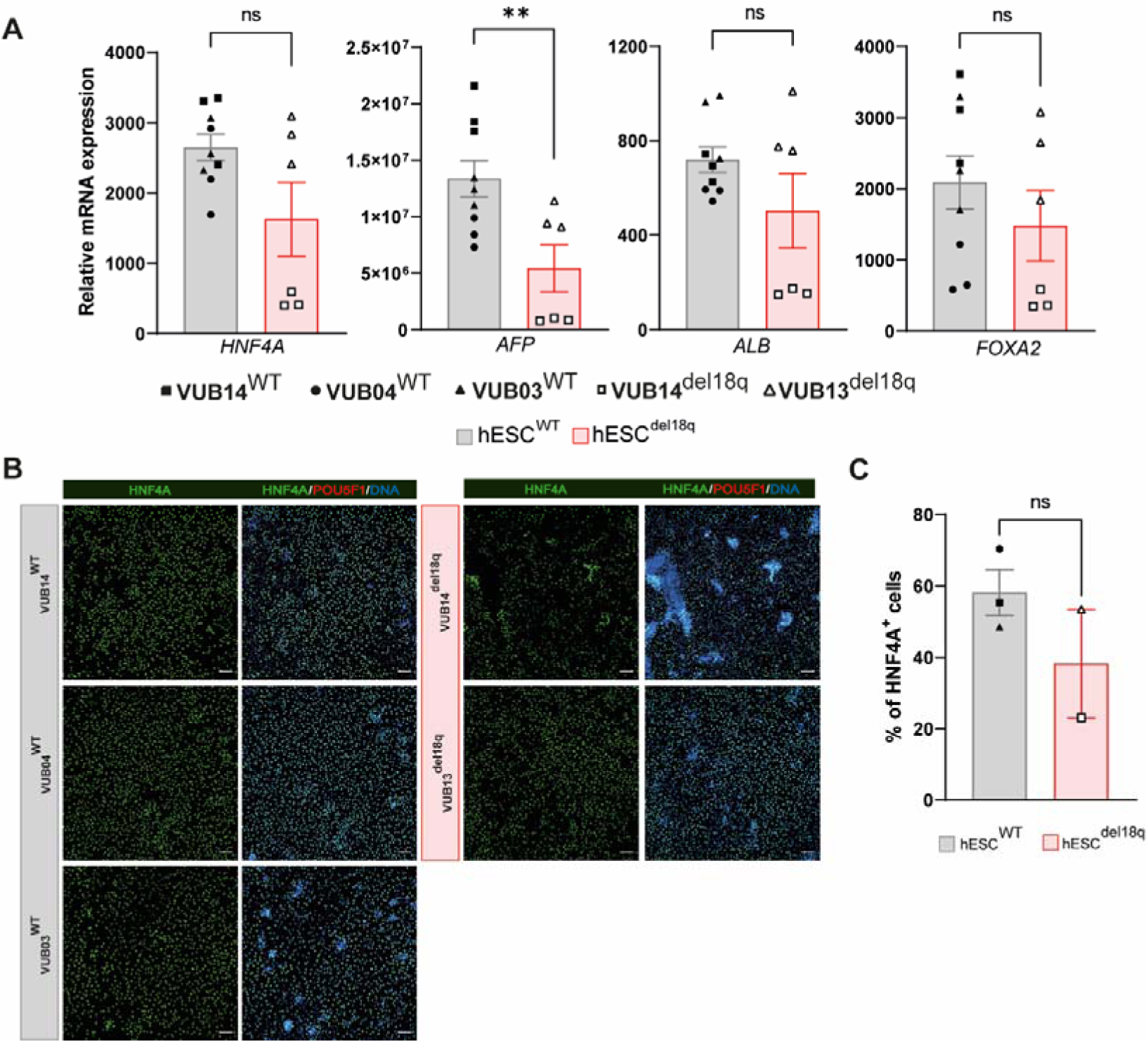
hESC^del18q^ and their genetically balanced counterparts differentiate equally well into hepatoblasts. (A) Relative mRNA expression of the cardiac progenitor markers *HNF4A*, *AFP*, *ALB* and *FOXA2*, as measured by qRT-PCR (n = 3). Data are shown as the means ± SEM; different patterns indicate different cell lines, and the horizontal bars with asterisks *, **, *** and **** represent statistical significance between samples at 5%, 1%, 0.1% and 0.01% respectively (unpaired t-test). (B) Immunostaining for HNF4A (green) and POU5F1 (red) in mutant and control lines after 8 days of hepatoblast differentiation, all scale bars are 100 µm. (C) Percentage of HNF4A-positive cells in the immunostainings shown in B.

For both mesodermal and endodermal lineage commitment, all hESC^WT^ and hESC^del18q^ lines displayed low mRNA and protein levels of undifferentiated state markers (Sup Fig. 2B-C), indicating a loss of pluripotency for all cell lines during differentiation. Overall, our results indicate that hESC^WT^ and hESC^del18q^ differentiate into definitive mesoderm and endoderm and that there may be a delay or impairment in the progression of hESC^del18q^ toward the cardiac progenitor stage. The differences in gene expression observed during hepatoblast differentiation are likely due to between-line variation in differentiation propensity rather than to the 18q deletion itself.

### Downregulation of *SALL3* impairs neuroectoderm differentiation but does not affect differentiation into mesoderm and endoderm

We next examined *SALL3* mRNA expression levels in undifferentiated cells and found that *SALL3* expression was significantly lower in hESC^del18q^ than in hESC^WT^ (Sup Fig. 3A), supporting the notion that *SALL3* could be a key gene in the altered differentiation capacity of hESCs^del18q^. We first generated *SALL3* knockdown (KD) lines from three hESC^WT^ lines (VUB02^WT^, VUB03^WT^ and VUB04^WT^) by transducing a lentiviral vector containing shRNA targeting the *SALL3* transcript (hESC^WT_*SALL3*KD^) or a nontargeting shRNA as a control (hESC^WT-NT^). We confirmed the knockdown efficiency of the generated hESC^WT^*SALL3*^KD^ lines by measuring the *SALL3* mRNA expression levels and found that they were reduced by 30%, 40% and 20% in VUB02^WT_*SALL3*KD^, VUB03^WT_*SALL3*KD^ and VUB04^WT_*SALL3*KD^ respectively, compared to the controls (Sup Fig. 3B). Next, we generated hESC^del18q^ with stable overexpression of *SALL3* (hESC^del18q_*SALL3*OE^) by transducing VUB13^del18q^ and VUB14^del18q^ with the *SALL3* lentiviral vector, and we verified the overexpression by measuring *SALL3* mRNA levels, which were significantly increased in VUB13^del18q_*SALL3*OE^ (3-fold) and VUB14^del18q_*SALL3*OE^ (5-fold) compared to controls (Sup Fig. 3C).

To investigate the role of *SALL3* in regulating hESC differentiation propensity, we first induced neuroectodermal differentiation in hESC^WT_*SALL3*KD^, hESC^del18q_*SALL3*OE^, and the corresponding control cells. All three hESC^WT_*SALL3*KD^ lines had lower levels of all NE markers than nontarget controls (Fig. 5A and Sup Fig. 4A-B). PAX6 protein levels were also reduced in hESC^WT_*SALL3*KD^ (Fig. 5B-C), with only 20% of the cells expressing PAX6, compared to 80% of PAX6 positive cells in the control group (Fig. 5B-C). Our results show that *SALL3* suppression in hESC^WT^ recapitulates the impaired NE differentiation seen in hESC^del18q^ lines. In contrast, hESC^del18q_*SALL3*OE^ cells efficiently differentiated into neuroectoderm cells, accompanied by a significant increase in the mRNA levels of *PAX6*, *SOX1*, and *NES* (Fig. 5A and Sup Fig. 4A-B); moreover, hESC^del18q*SALL3*OE^ cultures included more PAX6 positive cells than hESC^del18q^ cultures (60% vs 20%, respectively) (Fig. 5B-C). These results indicate that exongenous *SALL3* expression can rescue the impairment of ectoderm differentiation caused by 18q loss.

**Figure 5.**
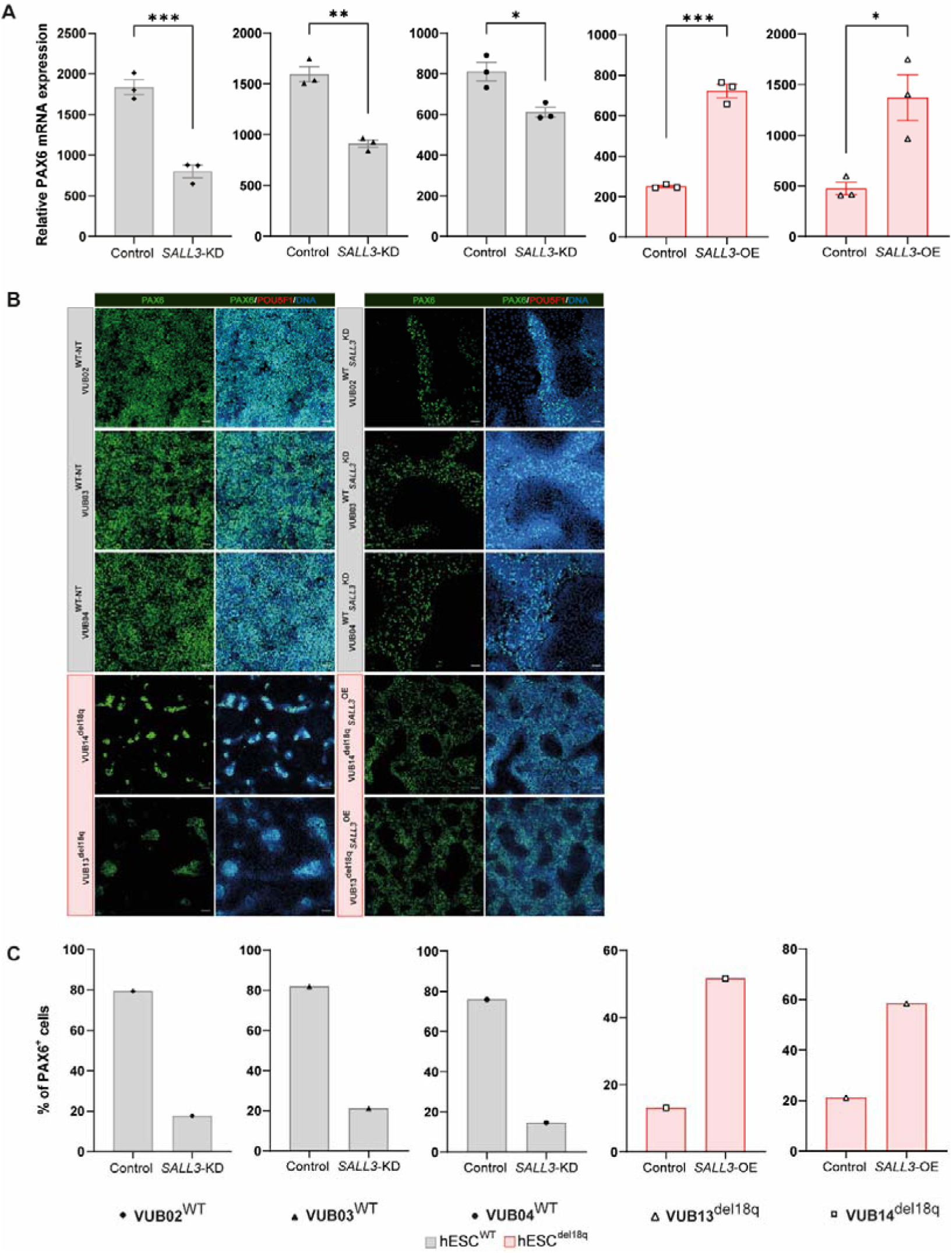
Downregulation of *SALL3* drives the impaired neuroectoderm differentiation of hESCs with 18q loss. (A) Relative mRNA expression as measured by qRT-PCR for the ectoderm marker *PAX6* (n = 3). Data are shown as the means ± SEM; different patterns indicate different cell lines, and the horizontal bars with asterisks *, **, *** and **** represent statistical significance between samples at 5%, 1%, 0.1% and 0.01% respectively (unpaired t-test). (B) Immunostaining for PAX6 (green) and POU5F1 (red) in mutant and control lines after 8 days of neuroectoderm differentiation, the scale bars in knock down groups are 50 µm and the scale bars in over-expression groups are 100 µm. (C) Percentage of PAX6-positive cells in the immunostainings shown in B.

Next, we induced the differentiation of the different hESC lines into cardiac progenitors. The three hESC^WT_*SALL3*KD^ lines differentiated inconsistently toward mesoderm fates. VUB04^WT_*SALL3*KD^ showed lower levels of *GATA4* (Fig. 6A, p=0.0003, unpaired t-test), whereas compared to the controls, VUB03^WT_*SALL3*KD^ and VUB02^WT_*SALL3*KD^ showed no differences in *GATA4* expression (Fig. 6A). Additionally, the percentage of GATA4^+^ cells detected by immunostaining in hESC^WT_*SALL3*KD^ followed the same pattern, consistent with the mRNA levels of each line (Fig. 6B-C). The hESC^del18q_*SALL3*OE^ cells exhibited variable marker profiles, with increases in GATA4 expression at both the mRNA (Fig. 6A, p=0.004, unpaired t-test) and protein levels for VUB13^del18q_*SALL3*OE^ but no difference in *GATA4* mRNA levels (Fig. 6A, p=0.3622, unpaired t-test) and a slight decrease in GATA4 protein levels in VUB14^del18q_*SALL3*OE^ (Fig. 6B-C). Similarly, the mRNA expression levels of other markers, *NKX2-5*, *ISL1*, and *PDGFRA,* showed no consistent trend in the hESC^WT_*SALL3*KD^ groups and exhibited no consistent differences in hESCs^del18q^ and hESC^del18q^*SALL3*^OE^ (Sup Fig. 5A-C).

**Figure 6.**
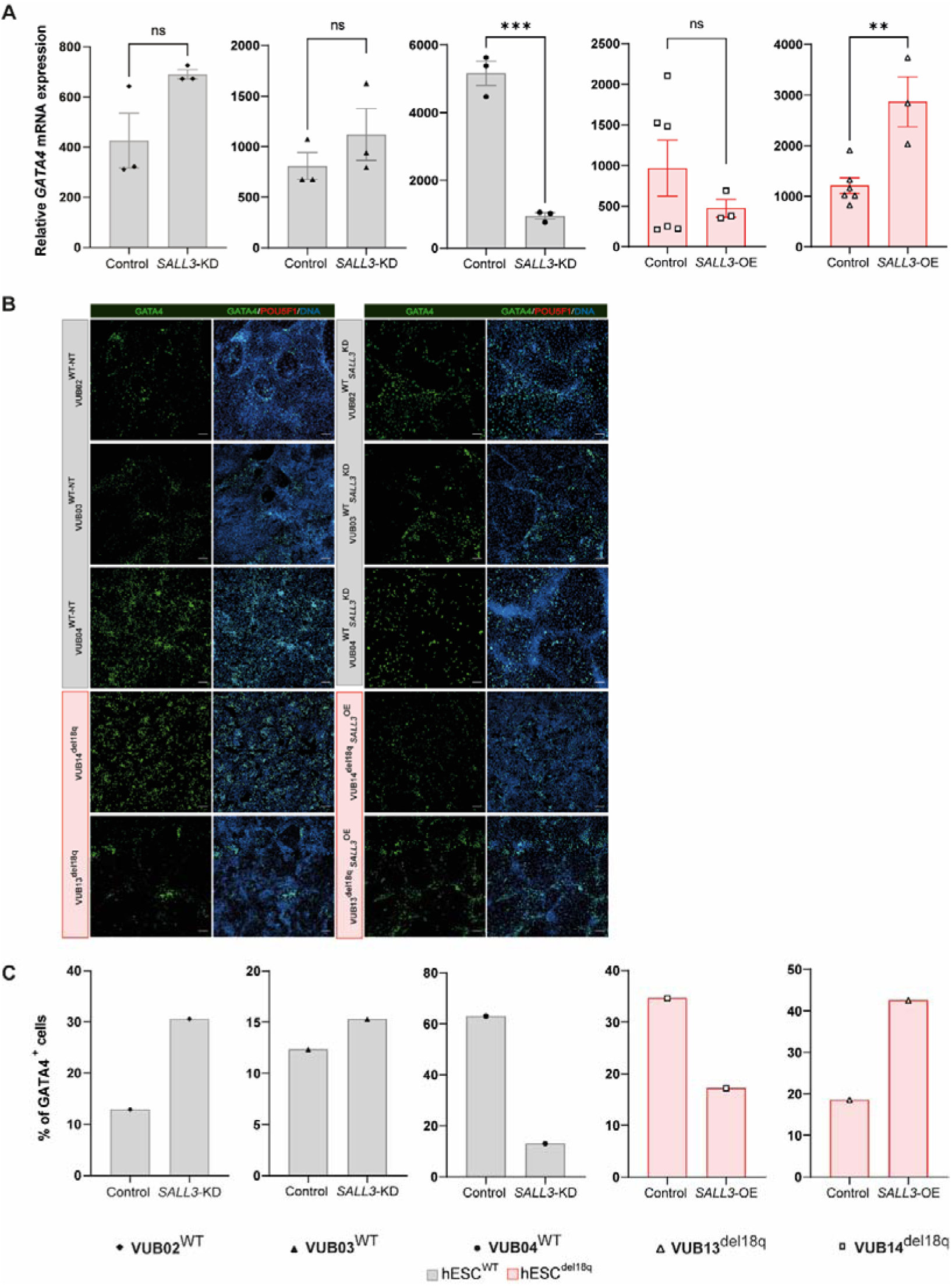
Changes in *SALL3* expression do not regulate cardiac progenitor differentiation. (A) Relative mRNA expression as measured by qRT-PCR for the cardiac progenitor marker *GATA4* (n = 3). Data are shown as the means ± SEM; different patterns indicate different cell lines, and the horizontal bars with asterisks *, **, *** and **** represent statistical significance between samples at 5%, 1%, 0.1% and 0.01% respectively (unpaired t-test). (B) Immunostaining for GATA4 (green) and POU5F1 (red) in mutant and control lines after 5 days of cardiac progenitor differentiation, all scale bars are 100 µm. (C) Percentage of GATA4-positive cells in the immunostainings shown in B.

When hESCs were differentiated into hepatoblast, *SALL3* downregulation in hESC^WT^ resulted in increased *HNF4A*, *ALB*, and *FOXA2* mRNA expression (Fig. 7A and Sup Fig. 4B-C). The percentage of HNF4A positive cells was higher in VUB03 ^WT^*SALL3*^KD^ cells but not in VUB04^WT_*SALL3*KD^ and VUB02^WT_S*ALL3*KD^ cells compared to controls (Fig. 7B-C). The mRNA expression of another marker, *AFP*, also did not show consistent changes (Sup Fig. 4A). Upon overexpression of *SALL3*, the differentiation profiles into hepatoblast did not show the expected mirroring effect. The mRNA expression of all the hepatoblast markers *HNF4A*, *ALB*, *AFP* and *FOXA2* increased in VUB14^del18q_*SALL3*OE^ cells, consistent with the changes observed in the HNF4A protein level, but this same effect was not observed in VUB13^del18q_*SALL3*OE^ cells (Fig. 7 and Sup Fig. 4).

**Figure 7.**
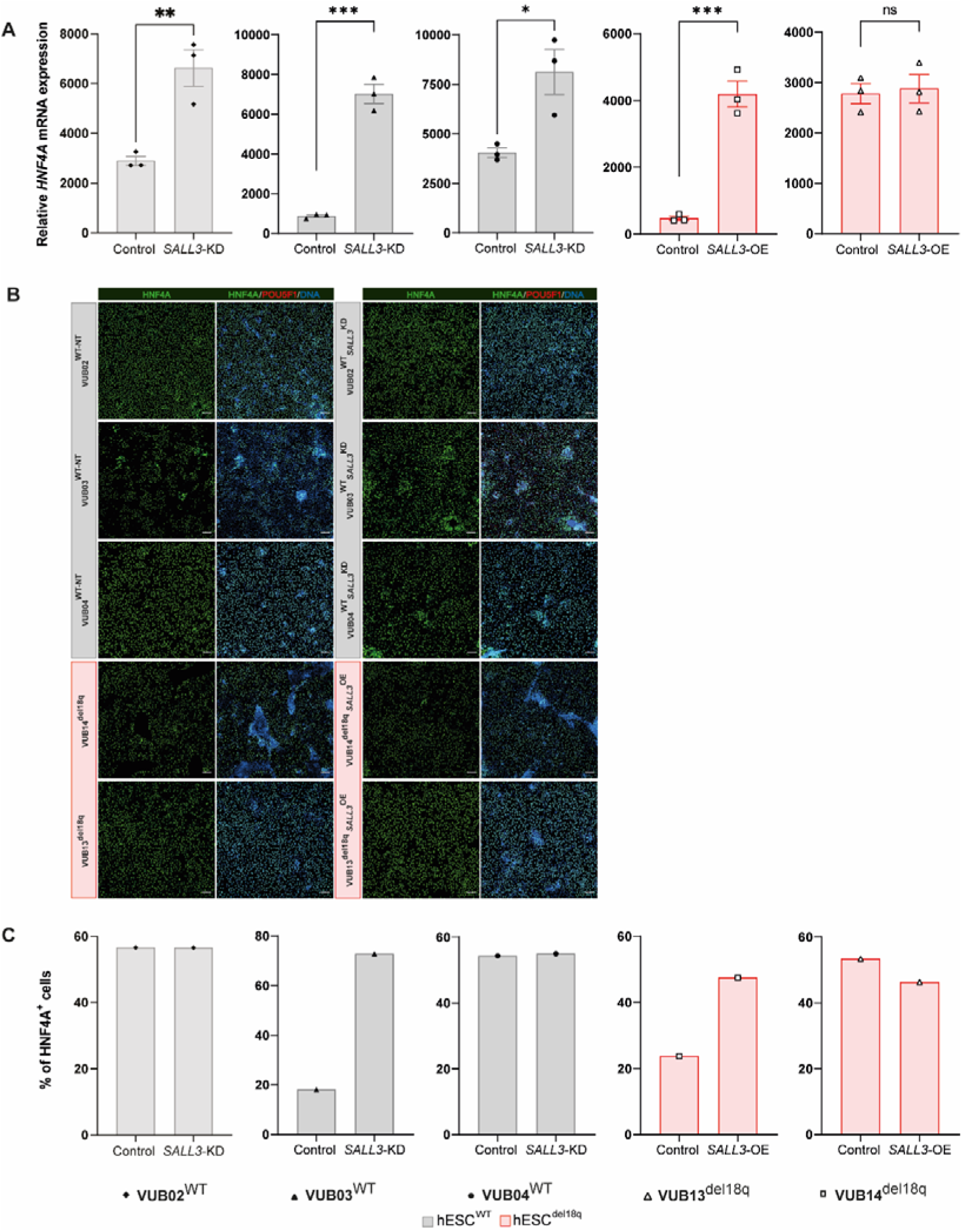
Changes in *SALL3* expression do not impact hepatoblast differentiation. (A) Relative mRNA expression of the ectoderm marker *HNF4A* as measured by qPCR (n = 3). Data are shown as the means ± SEM; different patterns indicate different cell lines, and the horizontal bars with asterisks *, **, *** and **** represent statistical significance between samples at 5%, 1%, 0.1% and 0.01% respectively (unpaired t-test). (B) Immunostaining for HNF4A (green) and POU5F1 (red) in mutant and control lines after 8 days of hepatoblast differentiation, all scale bars are 100 µm. (C) Percentage of HNF4A positive cells.

Overall, these results suggest that the effect of *SALL3* on mesoderm and endoderm differentiation may be line-specific and is not the reason for the delayed progression seen during cardiac differentiation in hESCs^18q^.

### Downregulation of *SALL3* and loss of 18q result in the deregulation of genes in pathways associated with pluripotency and differentiation

To gain deeper insight into the effect of *SALL3* downregulation on the global transcriptomic profile of cells with 18q loss, we carried out bulk RNA sequencing of hESC^WT-NT^ (N=6), hESC^WT_*SALL3*KD^ (N=6), hESC^WT^ (N=6), VUB13^del18q^ (N=4) and VUB13^del18q_*SALL3*OE^ (N=5) cells. To estimate the similarity among the samples, we generated a distance clustering heatmap with global row scaling (Fig. 8A) and performed principal component analysis (Fig. 8B). The heatmap shows that while VUB13^del18q^ and hESC^WT_*SALL3*KD^ cluster together, they cluster apart from the WT cell lines, as well as from VUB13^del18q_*SALL3*OE^. The samples in this last group clustered more closely to the WT cell lines than to their unmodified VUB13^del18q^ (Fig. 8A). This pattern was also reflected in the first dimension of the PCA (Fig. 8B), where hESC^WT^, hESC^WT-NT^ and VUB13^del18q_*SALL3*OE^ clustered more closely with each other than with VUB13^del18q^ and hESC^WT_*SALL3*KD^ (Sup Fig. 6A). These results suggest that the downregulation of *SALL3* in WT cells is sufficient to alter the transcriptome such that its profile is closer to that of an hESC line with a loss of 18q, and *SALL3* overexpression in hESC^del18q^ can restore the transcriptome to a near-WT state. Taken together, these findings support the our hypothesis that the differences between hESC^WT^ and hESC^del18q^ are mostly driven by the downregulation of *SALL3* due to the loss of one copy of this gene. For this reason, we pooled the hESCs^WT-NT^ with the untreated hESCs^WT^ for further analysis.

**Figure 8.**
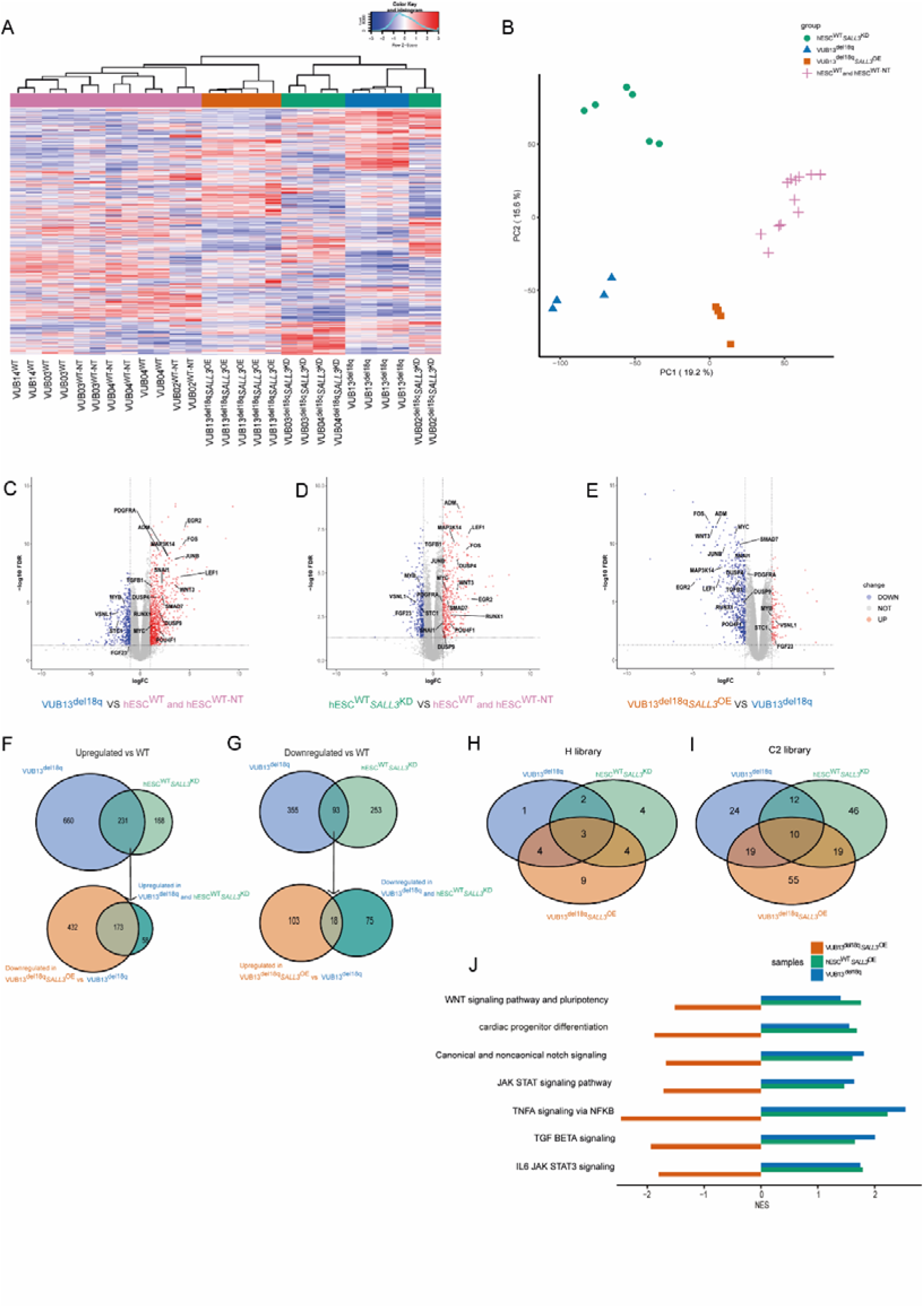
Downregulation of *SALL3* and loss of 18q result in the deregulation of genes in pathways associated with pluripotency and differentiation. (A,B) Unsupervised clustering heatmap(A) and Principal component analysis (PCA) (B) of the coding genes with a count per million greater than one in at least two samples. (C, D, E) Volcano plots of differential gene expression analysis for VUB13^del18q^ versus hESC^WT^ and hESC^WT-NT^(C), hESC^WT_*SALL3*KD^ versus hESC^WT^ and hESC^WT-NT^(D), and VUB13^del18q_*SALL3*OE^ versus VUB13^del18q^ (E) with a cutoff value of |log_2_fold change|> 1 and FDR < 0.05. (F) Venn diagrams showing the genes downregulated in VUB13^del18q_*SALL3*OE^ and upregulated in VUB13^del18q^ and hESC^WT_*SALL3*KD^. (G) Venn diagrams showing the genes upregulated in VUB13^del18q_*SALL3*OE^ and downregulated in VUB13^del18q^ and hESC^WT_*SALL3*KD^. (H) Venn diagrams of the pathways in the H library common among VUB13^del18q^, hESC^WT_*SALL3*KD^, VUB13^del18q_*SALL3*OE^ and hESC^WT^. (I) Venn diagrams of the pathways in the C2 library common among VUB13^del18q^, hESC^WT_*SALL3*KD^, VUB13^del18q_*SALL3*OE^ and hESC^WT^. (J) Pathways commonly deregulated among the different samples are associated with pluripotency and differentiation.

Next, we carried out differential gene expression analysis to gain further insight into which genes and pathways form the basis of the differences between hESC^WT^ and hESC^del18q^ and which of these are driven by *SALL3*. Figure 8B-D shows volcano plots of the differentially expressed genes in the different groups. We considered this differential expression to be significant at a |log_2_fold-change| > 1.0 and false discovery rate (FDR)< 0.05. VUB13^del18q^ shows 891 and 448 genes that are significantly upregulated and downregulated in hESC^del18q^, respectively, compared to the WT cells (Fig 8C). Downregulation of *SALL3* in WT cells led to the differential expression of 745 genes, of which 399 were upregulated and 346 were downregulated (Fig 8D). Overexpression of *SALL3* in VUB13^del18q^ resulted in the upregulation of 121 genes and the downregulation of 605 genes (Fig 8E).

To elucidate which of the transcriptional differences between hESC^WT^ and hESC^del18q^ are mediated by *SALL3*, we investigated the overlaps in up- and downregulated genes across conditions. First, we found that 231 and 93 genes were commonly up- and downregulated, respectively, between VUB13^del18q^ and hESC^WT_*SALL3*KD^, representing 26% of the differentially expressed genes in hESCs with an 18q deletion (Fig 8F and G). Next, we compared these subsets of genes to the genes with altered gene expression in hESC^del18q^ upon overexpression of *SALL3*. We compared the genes that were upregulated by *SALL3* knockdown and by loss of 18q to those downregulated by overexpression of *SALL3* in hESC^18q^, and vice versa (Fig 8F and G). Out of the 231 upregulated genes shared by the VUB13^del18q^ and hESC^WT_*SALL3*KD^ groups, 173 genes showed increased expression in VUB13^del18q_*SALL3*OE^ (Fig 8F). Additionally, 18 of the 93 downregulated genes shared by VUB13^del18q^ and hESC^WT_*SALL3*KD^ displayed increased expression in VUB13^del18q^*SALL3*^OE^ (Fig 8G). This further refined the gene set to a core of 191 genes that are the most strongly regulated by *SALL3* in hESCs, both by the loss of a copy of the gene itself and by the modulation of its expression (Supplementary Table 2). The other differences in gene expression are likely associated with the loss of other genes in 18q or with the overexpression of genes in the duplicated regions of chromosomes 5 and 7, which are part of the derivative chromosome 18 in hESC^del18q^.

Finally, to identify potential molecular targets and elucidate the underlying functional mechanisms that contribute to the observed impairment of differentiation capacity in hESC^del18q^, we analyzed the differential gene expression of the three groups using Gene Set Enrichment Analysis (GSEA) and the MSigDB database. Specifically, we focused on the Kyoto Encyclopedia of Genes and Genomes (KEGG) and WikiPathways databases within the C2 library and the pathways of the H library. We filtered the significant pathways based on a normalized enrichment score |NES| > 1, a p-value < 0.05, and the proportion of leading-edge genes accounting for over 30% of the entire gene set involved in the pathway.

Figures 8H and I show Venn diagrams of the overlap between significantly enriched pathways in the C2 and H libraries, respectively, for each of the three groups. The full list can be found in the Supplementary Table 3. In total, we found 13 pathways that overlapped among the three groups, all 13 of which were positively enriched in both VUB13^del18q^ and hESC^WT_*SALL3*KD^ and negatively enriched in VUB13^del18q_*SALL3*OE^ (Fig 8J shows the cell-type relevant pathways; the whole set can be found in the Supplementary Table 4). Because of the critical role that these pathways play in pluripotency maintenance and differentiation, we analyzed the expression of pluripotency-associated genes, and we found that *NANOG, POU5F1, LIN28, SOX2, PODXL, SUSD2, MYC, FOXD3* and *DPPA3* are overexpressed in VUB13^del18q^, with all but *DPPA3* and *POU5F1* also being overexpressed upon *SALL3*^KD^ and downregulated by transgenic *SALL3* overexpression in VUB13^del18q^ (Sup Fig. 7A).

## Discussion

In this study, we examined the repercussions of the loss of chromosome 18q on the differentiation capacity of hESCs. For this, we used an early-stage differentiation approach to generate lineage-specific cell types representing the three germ layers: neuroectoderm, hepatoblast and cardiac progenitors. Our *in vitro* lineage commitment studies indicated that the deletion of 18q in hESCs impaired neuroectodermal differentiation and delayed cardiac progenitor differentiation, while no consistent differences were observed in the commitment toward hepatoblasts. To study the mechanistic basis for these changes in differentiation, we looked at the genes located in the minimal region of loss. We found that decreased *SALL3* expression due to the loss of one copy of the gene was sufficient to result in the observed decreased neuroectoderm differentiation but not to modulate cardiac and hepatoblast differentiation. In this sense, our results are only partially aligned with those obtained by Kuroda *et al.*^51^. These authors found that downregulating *SALL3* resulted not only in decreased neuroectoderm differentiation, similar to our results, but also in increased cardiac progenitor differentiation. While we can only speculate about the reasons for these differences, it is likely that the genetic background of the cell lines plays an important role^59^. Also, given that *SALL3* is a modulator of DNMT3A activity^60^, the preexisting epigenetic marks in each of the lines, particularly histone modifications, may influence the recruitment of DNMT3A to methylate the DNA^61, 62^. Furthermore, it is possible that other genes located in 18q, or in the gain regions of chromosomes 5 and 7, cause cell-line specific effects.

In line with this reasoning, the gene expression analysis revealed that only part of the divergence between hESCs with 18q loss and their chromosomally normal isogenic counterparts was related to the differential expression of *SALL3*. Interestingly, the core set of deregulated genes were associated with pathways involved in maintenance of and exit from the undifferentiated pluripotent state, and key regulators of pluripotency were both upregulated in hESCs^18q^ and regulated by *SALL3* expression. For instance, TGF-beta signaling is key to the maintenance of the primed pluripotent state^63^. TGF-beta and BMP4 signaling are also core genes in regulating the balance between neuroectoderm differentiation and mesendoderm in both humans and mice^64^. Notch activation mediates TGF-β signaling during hESC and mesenchymal stem cell differentiation into smooth muscle cells^65^, and its inhibition supports naïve state consolidation in rodent models^63^. The cytokine tumor necrosis factor alpha has been shown to negatively regulate the differentiation of various cell types, including cardiomyocytes^66^, embryoid bodies^67^ and osteoblasts^68, 69^. Additionally, nuclear factor-κB inhibition has been found to mediate naïve pluripotency in mice^70^. Overall, these results suggest that downregulation of *SALL3* due to the loss of 18q alters undifferentiated-state maintenance in hESCs by affecting pluripotency-associated pathways, and these changes have profound effects on the differentiation capacity of the cells.

With hPSCs steadily moving into clinical trials^71^ and being broadly used as a cell source for *in vitro* modelling of, for instance, developmental processes and diseases, determining the impact of recurrent genetic abnormalities is critical^72–74^. Work from our group and others is beginning to generate a detailed picture showing how these genetic abnormalities affect differentiation in a cell lineage-specific manner. For instance, 20q11.21 gain impairs neuroectoderm commitment without affecting mesendoderm induction^75, 76^, and recently, it has been shown that cells with an isochromosome 20q are not able to survive RPE differentiation and display overall disruptions in the ability to correctly differentiate^77^. In this work, we show that 18q loss specifically impairs neuroectoderm commitment and appears to delay cardiac differentiation. Taken together, these results highlight the importance of the genetic screening of hPSC cultures to ensure that these abnormalities do not pass unnoticed. In a research setting, chromosomal abnormalities could lead to confounding effects that decrease the reliability and reproducibility of the work, and in a clinical setting, they could lead at best to decreased therapeutic efficacy and at worst to tumorigenesis^72–74,78,79^.

Losses of chromosome 18q are recurrently found in cancers ^33^ and have been shown to be early events in midgut carcinoids^80, 81^. Loss of heterozygosity on chromosome 18q is associated with significantly decreased survival in head and neck squamous cell carcinoma patients, and the methylation status of the *SALL3* promoter correlates with shortened disease-free survival^82–84^. Furthermore, three cancer suppressor genes have been identified in 18q, PIGN, MEX3C and ZNF516, which play roles in replication stress and in cervical neoplasia^85–87^.

In this context, several studies have found that even when chromosomally abnormal hPSCs are capable of differentiating, they display altered gene expression patterns suggestive of malignant transformation^72, 88–91^. Furthermore, the abnormalities seen in hPSCs could be regarded as a first hit in cancerous transformation. Cells that are already genetically abnormal upon transplantation are not cancerous yet but may have a higher chance of undergoing oncogenic transformation as they could require fewer additional genetic hits to initiate the process^74, 78, 79^. With this in mind, centers involved in clinical work subject their hPSC cultures and hPSC-based products to genetic screening prior to their use in patients. This typically involves the use of G-banding and, increasingly commonly, massively parallel sequencing (MPS), which enable the detection of chromosomal abnormalities and even, in the case of MPS, potentially harmful single nucleotide changes. Incidentally, the first clinical trial using hPSC-derived retinal pigmented epithelium (RPE) was halted after potentially harmful mutations were identified in both the hPSCs and the RPE cells derived from this line^92^. Unfortunately, standard methods for genetic screening cannot detect abnormalities present as low-grade mosaics in the hPSC culture, which are known to be common^93–95^. If any of the cells carry an abnormality that specifically impairs the differentiation to the chosen lineage, their presence could lead to a cell product containing a subpopulation of poorly differentiated or mis-specified cells and, in the worst-case scenario, with tumor-initiating capacity. This risk further highlights the need for more detailed knowledge of the impacts of specific abnormalities on the properties of hPSCs so that targeted screening approaches that can detect potentially small populations of abnormal cells in cultures can be developed.

In conclusion, in this study, we have characterized the differentiation capacity of hESCs with 18q loss, one of the recurrent, albeit less common, genetic abnormalities found in hPSC cultures. We found that these cells are characterized by abnormal differentiation into cardiac progenitors and an impaired capacity for neuroectoderm commitment, the latter driven by the loss of one copy of *SALL3*. This gene is an inhibitor of *DNMT3A*, and its downregulation results in changes in the expression of genes involved in the maintenance of pluripotency and in hESC differentiation. Further research will be needed to assess whether other cell-type specific effects of this abnormality exist and might be revealed by longer differentiation protocols, beyond the progenitor stage, as well as the potential consequences of 18q loss for oncogenic potential.

## Materials and methods

### hESC maintenance and passaging

All hESC lines were derived and characterized as reported previously^96, 97^ and are also registered in the EU hPSC registry (https://hpscreg.eu/). They were cryopreserved in freezing medium composed of 90% knock-out serum (Thermo Fisher Scientific) mixed with 10% dimethyl sulfoxide (Sigma-Aldrich). The hESCs were maintained in NutriStem hESC XF medium (NS medium; Biological Industries) with 100 U/mL penicillin/streptomycin (P/S) (Thermo Fisher Scientific) in a 37 °C incubator with 5% CO_2,_ and the culture medium was changed daily. The tissue culture dishes and plates (Thermo Scientific) were coated with 10 µg/mL Biolaminin 521 (Biolamina®) at 4 °C and then incubated at 37 °C for at least 20 min before the cells were seeded. The cells were passaged as single cells using TrypLE Express (Thermo Fisher Scientific) and split at a ratio of 1:10 to 1:100 as needed at 70–90% confluence. The medium was supplemented with 10 μM Rho kinase (ROCK) inhibitor Y-27632 (ROCKi, Tocris) for the first 24 h after passaging.

### Copy number variant (CNV) analysis

The genetic content of the hESCs was assessed through shallow whole-genome sequencing by the BRIGHTcore of UZ Brussels, Belgium, as previously described^98^. We also conducted copy number variant analysis using quantitative real-time PCR (qRT-PCR) at regular intervals, particularly before and after performing lentiviral transduction and starting differentiation. DNA was extracted with a DNeasy Blood and Tissue Kit (Qiagen) according to the manufacturers’ protocol. qPCR was performed with the subsequent copy number assays: *RNaseP* (Thermo Fisher Scientific) as a reference and *KIF14*, *NANOG*, *NMT1*, and *ID1* (Thermo Scientific) representing the 1q, 12p, 17q and 20q regions, respectively, which are commonly subject to CNV in hESCs. The reaction systems were prepared by mixing *RNaseP*, TaqMan 2× Mastermix Plus – Low ROX (Eurogentec), and the related TaqMan assays together with 8 µl of the diluted DNA samples (n=3). qPCR was performed on a ViiA7 thermocycler (Thermo Fisher Scientific), and Applied Biosystems Copy Caller v.2.1 was used to analyze the CNVs.

### Total RNA isolation and cDNA synthesis

Total RNA was isolated using RNeasy Mini and Micro kits (Qiagen) following the manufacturer’s guidelines, including on-column DNase I treatment. The isolated RNAs were collected in nuclease-free water and analyzed for quality and quantity using UV spectrophotometry. The final RNA was stored at −80 °C. A minimum of 500 ng of mRNA was reverse-transcribed into biotinylated cDNA using the First-Strand cDNA Synthesis Kit (Cytiva) with the NotI-d(T)18 primer, and the resulting cDNA was stored at -20 °C for subsequent analysis.

### Quantitative real-time PCR (qRT-PCR) for gene expression analysis

Quantitative real-time PCR (qRT-PCR) was carried out using TaqMan mRNA expression assays (Thermo Fisher Scientific) and TaqMan 2× Mastermix Plus – Low ROX (Eurogentec) on a ViiA 7 thermocycler (Thermo Fisher Scientific) using the standard cycling protocol provided by the manufacturer. The relative expression of target genes was quantified using the comparative threshold cycle (Ct) method and normalized to the TaqMan GUSB transcript (Applied Biosystems) as the endogenous housekeeping gene. All the samples were run in triplicate, and the related TaqMan assays used in the present study are listed in Supplementary Table 5.

### Immunostaining

Differentiated cells were first fixed in a solution of PBS containing 3.7% formaldehyde (Sigma-Aldrich) for 15 min, permeabilized in 0.1% Triton X for 10 min (Sigma-Aldrich) and then blocked with 10% fetal bovine serum (ThermoFisher Scientific) for 1 h at room temperature (RT). Sequentially, primary antibodies appropriately diluted in blocking solution (1:200 dilution in 10% FBS) were incubated overnight at 4 °C. Thereafter, secondary antibodies conjugated to Alexa 488, Alexa 594 or Alexa 647 (1:200 dilutions in 10% FBS, Thermo Fisher Scientific) and Hoescht (1:1000 dilution, ThermoFisher Scientific) were applied for 1-2 h at room temperature in the dark. PBS was used to wash the cells three times between steps. Confocal images were acquired with an LSM800 confocal microscope (Carl Zeiss) with a 10× or 20× objective. For quantification, the fixed differentiated cells stained with respective antibodies were counted and compared to the number of Hoescht-stained nuclei to determine the percent positivity using Zen 2 (blue edition) imaging software. The areas are randomly selected (n=3-6) within a single well based on the Hoescht channel, and the positive cells are quantified by calculating the ratio of total positive cells to the total number of cells selected within those areas. The lists with antibodies can be found in supplementary Table 6.

### In vitro differentiation of iPSCs

#### Definitive neuroectoderm specification

The protocol for inducing neuroectoderm differentiation was adapted from the protocol described by Douvaras and colleagues^99^. In brief, hESCs were seeded on Biolaminin 521 (Biolamina®)-coated 24-well plates at a ratio of 100,000 cells per cm^2^ and grown to 90% confluence. Then, differentiation was induced by incubating the hESCs with neural induction specification medium for up to 8 days with daily medium changes. The neural induction specification medium consisted of basal medium supplemented with the following differentiation factors: 100 nM retinoic acid (Sigma-Aldrich), 10 µM SB431542 (Tocris), and 250 nM LDN193189 (STEMCELL Technologies). The basal medium was prepared by mixing DMEM/F12 (Thermo Scientific) basic medium supplemented with 1x NEAA (Thermo Scientific), 1x GlutaMAX (Thermo Scientific), 1x 2-mercaptoethanol (Thermo Scientific), 25 µg/mL insulin (Sigma-Aldrich) and 1x penicillin/streptomycin (P/S) (Thermo Scientific).

#### Cardiac progenitor differentiation

The induction of cardiac progenitor differentiation was initiated using a slightly modified version of the protocol from a previous publication^100^. hESCs were seeded on Biolaminin-521-coated 24-well plates at a density of 100,000 cells/cm^2^ and allowed to reach 80–90% confluence. At this point, differentiation was initiated by treating the hESCs with cardiomyocyte differentiation basal medium (CDBM) with 5 mM CHIR99021 (Tocris) to activate Wnt/β-catenin signaling and form the mesendoderm layer. After 24 h, CHIR99021 was removed, and the cells were cultured in CDBM with 0.6 U/mL heparin (Sigma-Aldrich) for 24 h. Subsequently, the CDBM medium was supplemented with 0.6 U/ml heparin and 3 mM IWP2 (Tocris) for another 3-day incubation with medium refreshed daily. The cardiomyocyte differentiation basal medium (CDBM) was composed of 10 µg/ml transferrin (Sigma Aldrich), 1x chemically defined lipid concentrate (Thermo Scientific) and E8 basal medium. The E8 medium was composed of DMEM/F12 (Thermo Scientific) basic medium with 64 mg/L L-ascorbic acid (Sigma-Aldrich) and 13.6 µg/L sodium selenium (Sigma-Aldrich).

#### Hepatoblast differentiation

Differentiation of hESCs into hepatoblasts was conducted using a protocol based on^101^. The hESCs were seeded at a density of 5x10^4^ cells/cm^2^ on 24-well plates precoated with Biolaminin-521. Upon reaching approximately 40-50% confluency, the hESCs were treated with liver differentiation medium (LDM) along with 50 ng/mL Activin A (STEMCELL Technologies), 50 ng/mL WNT3A (PeproTech) and 6 µL/mL DMSO (Sigma-Aldrich) for 48 h. The cells were incubated for an additional 48 h in the same medium without WNT3A. Then, the medium was changed to LDM with 50 ng/mL BMP4 (STEMCELL Technologies) and 6 µL/mL DMSO for the following 4 days, with the medium changed every two days. The samples were collected on day 8. LDM medium was created by adding MCDB 201 medium (pH=7.2, Sigma-Aldrich), L-ascorbic acid (Sigma-Aldrich), insulin-transferrin-selenium (ITS-G, Thermo Scientific), linoleic acid-albumin from bovine serum albumin (LA-BSA, Sigma-Aldrich), 2-mercaptoethanol (Thermo scientific), and dexamethasone (Sigma-Aldrich) to DMEM high glucose medium (Westburg Life Sciences).

### Generation of *SALL3* knock-down and overexpression cell lines

hESC^WT_*SALL3*KD^ cells were generated by infecting hESC^WT^ with lentiviral particles expressing *SALL3*-targeted shRNAs. To generate lentiviral particles, we transfected HEK 293T cells with individual clones from a SigmaMISSION shRNA targeting set (TRCN0000019754, TRCN0000417790) or control shRNA plasmid along with packaging plasmids (plasmid pMDG, encoding VSV G, and plasmid pCMVΔR8.9, encoding gag-pol). hESC^del18q_*SALL3*OE^ cells were generated by infecting hESC^del18q^ with lentiviral particles expressing *SALL3*. Lentiviral particles were generated as described above. The pLVSIN-EF1α puromycin vector expressing *SALL3* was a gift from Yoji Sato from the Division of Cell-Based Therapeutic Products, National Institute of Health Sciences, Japan. For transduction, hESCs were seeded at a density of 50,000/cm^2^ and then transduced with a 2:1 mix of NutriStem and lentivirus-containing medium in the presence of 10 mg/mL protamine sulfate (LEO Pharma) for 4 h. Cells were then washed with PBS before adding fresh NutriStem medium. Twenty-four hours later, the cells were again washed with PBS and then selected by puromycin.

### RNA Sequencing

RNA-seq library preparation was performed using QuantSeq 3′ mRNA-Seq Library Prep Kits (Lexogen) following Illumina protocols. Sequencing was performed on a high-throughput Illumina NextSeq 500 flow cell. On average, 13.9 x 10^6^ ± 7.1 x 10^6^ paired-end reads per sample were uniquely mapped, with an average coverage of 101 paired reads. The FastQC algorithm^102^ was used to perform quality control on the raw sequence reads prior to the downstream analysis. The raw reads were aligned to the new version of the human Ensembl reference genome (GRCh38.p13) with Ensembl (GRCh38.83gtf) annotation using STAR version 2.5.3 in 2-pass mode^103^.The aligned reads were then quantified, and transcript abundances were estimated using RNA-seq by expectation maximization (RSEM, version 1.3.3)^104^.

The count matrices were imported into R software (version 3.3.2) for further processing. The edgeR^105^ package was utilized to identify differentially expressed genes (DEGs) between groups. Transcripts with a count per million (cpm) greater than 1 in at least two samples were considered for the downstream analysis. Genes with a log2-fold change greater than 1 or less than -1 and a false discovery rate (FDR)-adjusted P-value less than 0.05 were considered significantly differentially expressed. Volcano plots of DEGs were generated using the ggplot2^106^ package in R, while Venn diagrams using VennDiagram function were used to visualize the overlap of the DEGs among different individuals.

Principal component analysis (PCA) and heatmap clustering were performed using normalized counts and R packages. The heatmap was generated using the heatmap.2 funtion. PCA was performed using the prcomp function and plotted using ggplot2. Gene set enrichment analysis (GSEA) was applied to detect the enrichment of pathways using the GSEA function in R with the MSigDB C2 and H databases. Values are ranked by sign(logFC)*(-log10(FDR)). |NES| > 1 and p-value < 0.05 were considered the thresholds for significance for the gene sets.

### Statistics

All differentiation experiments were carried out in at least triplicate (n ≥ 3). All data are presented as the mean ± standard error of the mean (SEM). Statistical evaluation of differences between 2 groups was performed using unpaired two-tailed t tests in GraphPad Prism9 software, with p < 0.05 determined to indicate significance.

### Data availability

The RNA sequencing counts per million tables are provided in the supplementary material. The raw sequencing data are available upon request.

## AUTHOR CONTRIBUTIONS

Y.L. carried out all of the experiments and bioinformatics analysis unless stated otherwise and co-wrote the manuscript. D.A. D. cowrote the manuscript. N.K. packaged the lentivirus and assisted in transduction. E.C.D.D. assisted with the bioinformatics analysis. M.R. assisted in microscopy and cell counting. C.J. and M. G assisted with the cell culture. K.S. proofread the paper. C.S. cowrote the manuscript and designed and supervised the experimental work.

## ACKNOWLEDGMENTS

The authors wish to thank Yoji Sato from the Division of Cell-Based Therapeutic Products, National Institute of Health Sciences in Japan for kindly sharing the lentiviral construct for *SALL3* overexpression. Y.L. is a predoctoral fellow supported by the China Scholarship Council (CSC), and M.R., C.J., N.K. and E.C.D.D. are predoctoral fellows supported by the Fonds voor Wetenschappelijk Onderzoek Vlaanderen (FWO). This research was supported by the FWO (grant number 1506617N) and the Methusalem Grant to Karen Sermon (Vrije Universitet Brussel).

