## supplemental data for "*SALL3* mediates the loss of neuroectodermal differentiation potential in human embryonic stem cells with chromosome 18q loss"

**Supplementary table 1. Passage range and karyotype of the lines in this study**

| Line | Passage range | Karyotype | Method of karyotyping and size in Mb |
| --- | --- | --- | --- |
| VUB14^wt^ | 51-55 | 46, XX | Shallow whole genome sequencing |
| VUB04^wt^ | 16-23 | 46, XX | Shallow whole genome sequencing |
| VUB03^wt^ | 19-25 | 46, XX | Shallow whole genome sequencing |
| VUB13^del18q^ | 52-58 | 46, XX  dup(5) (q21.3qter)  del(18) (q21.2qter) | Shallow whole genome sequencing  chr5: 33 Mb  chr18: 29 Mb |
| VUB14^del18q^ | 49-54 | 46, XX,  dup(7)(p22.3pter) del(18)(q21.32qter) | Shallow whole genome sequencing  chr7: 32 Mb  chr18: 21 Mb |
| VUB04 | 29 | 46, XX | Array-based comparative genomic hybridization (published in Spits et al., 2008) |
|  | 66 | 46, XX, dup(5)(q14.2qter), del(18)(q21.2qter) | Array-based comparative genomic hybridization (published in Spits et al., 2008)  Chr5: 99.5 Mb  Chr18: 28.7 Mb |
| VUB26 | 10 | 46, XX, dup(7)(q33qter), del(18)(q23qter) | Array-based comparative genomic hybridization (published in Spits et al., 2008)  Chr7: 11.3 Mb  Chr18: 4.6 Mb |

Supplementary figure 1. Shallow sequencing results of the lines used in this study

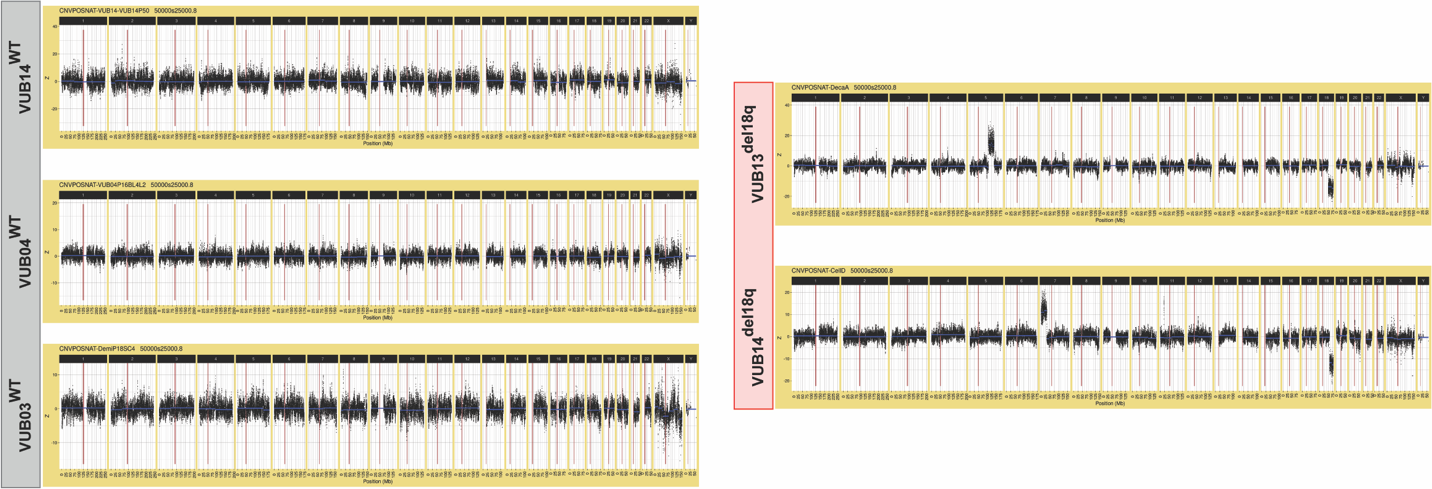

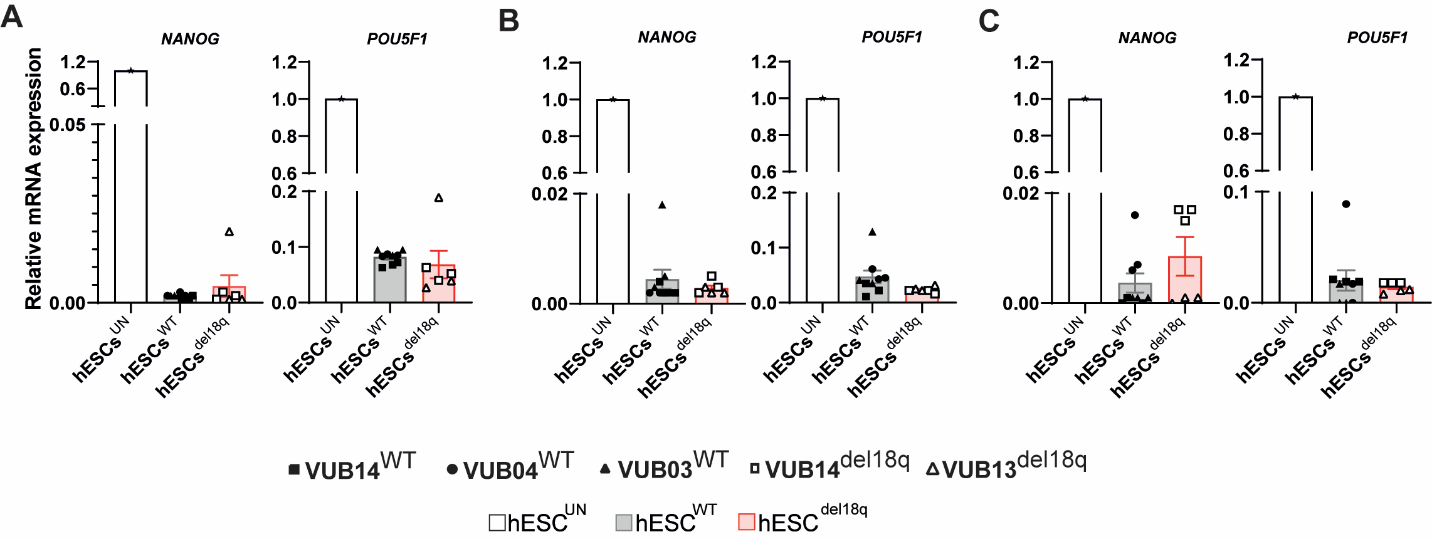

**Supplementary figure 2. The relative mRNA expression of undifferentiated state markers during the differentiation.** (A-C) Relative mRNA expression as measured by qRT-PCR for pluripotency markers *NANOG*, *POU5F1* during the differentiation process (n = 3). Data are shown as means ± SEM, each pattern indicates different cell lines. Neuroectoderm (A) cardiac progenitor (B) hepatoblasts (C).

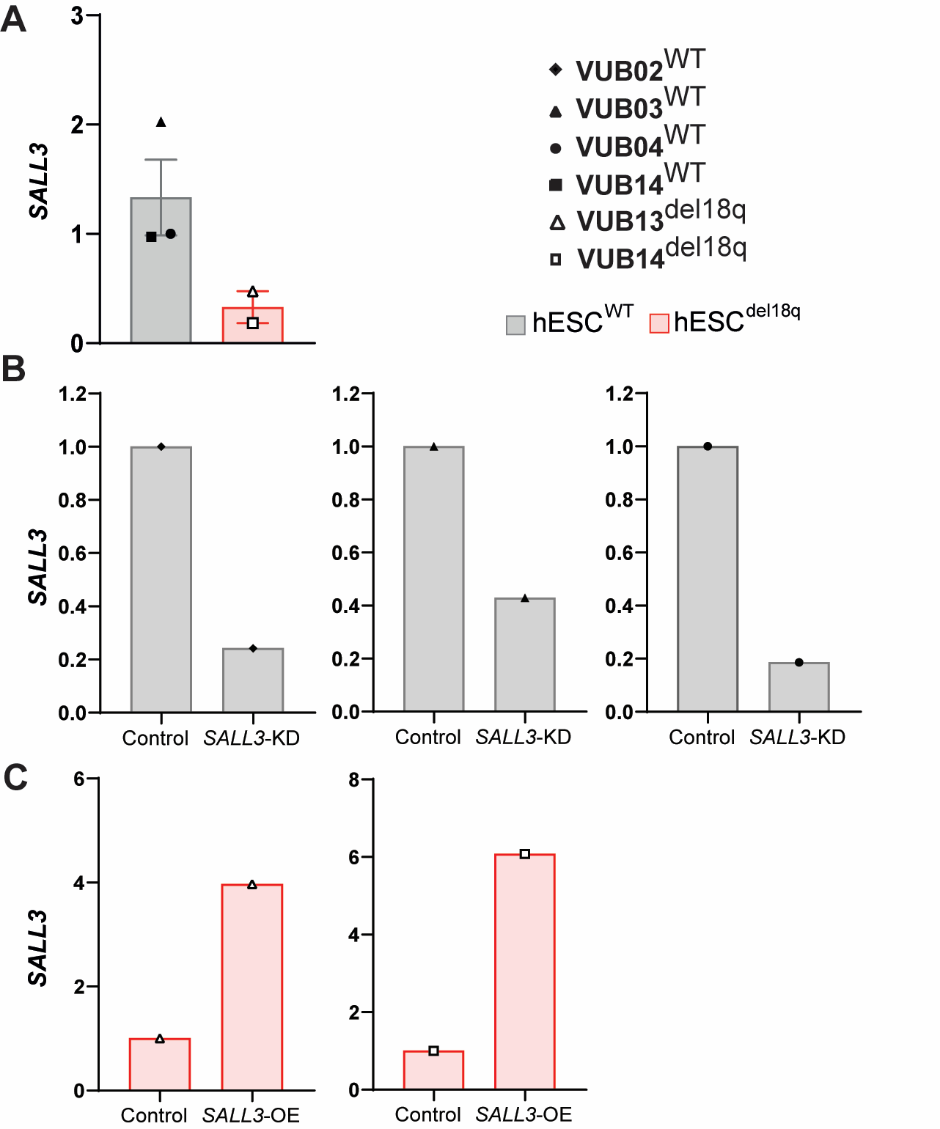

**Supplementary figure 3. The relative *SALL3* mRNA expression** (A) Relative *SALL3* mRNA expression as measured by qRT-PCR between hESC^del18q^ compared to hESC^WT^. (B) Relative *SALL3* mRNA expression as measured by qRT-PCR between hESC^WT-NT^ and hESC^WT_^*^SALL3^*^KD^. (C) Relative *SALL3* mRNA expression as measured by qRT-PCR between hESC^del18q^ and hESC^del18q_^*^SALL3^*^OE^.

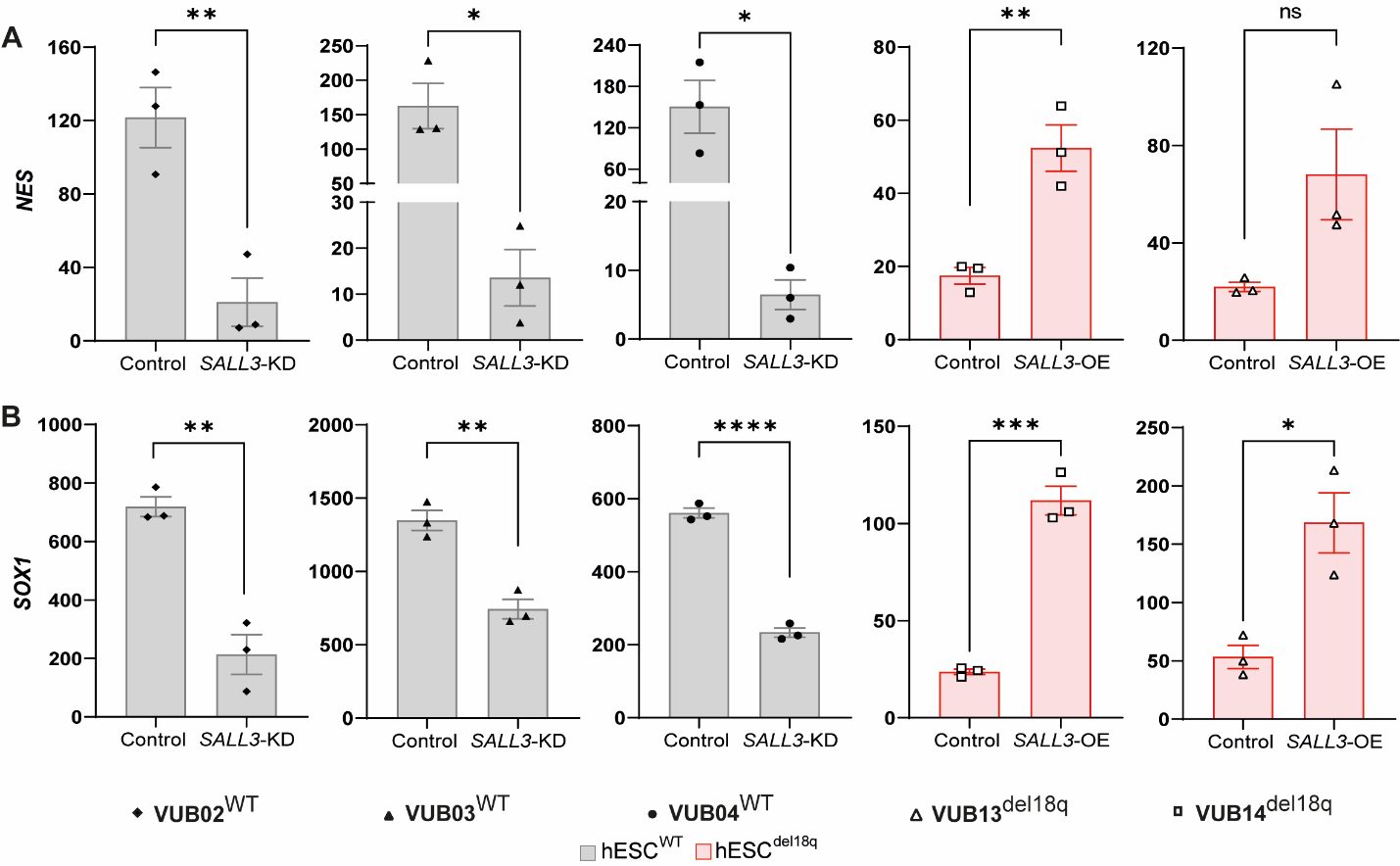

**Supplementary Figure 4. *SALL3* drives the impaired neuroectoderm differentiation** (A-C) Relative mRNA expression for neuroectoderm markers (n = 3) by qRT-PCR. Different patterns indicate different cell lines, and the horizontal bars with asterisks *, **, *** and **** represent statistical significance between samples at 5%, 1%, 0.1% and 0.01% respectively (unpaired t-test). *NES* (A), *SOX1*(B).

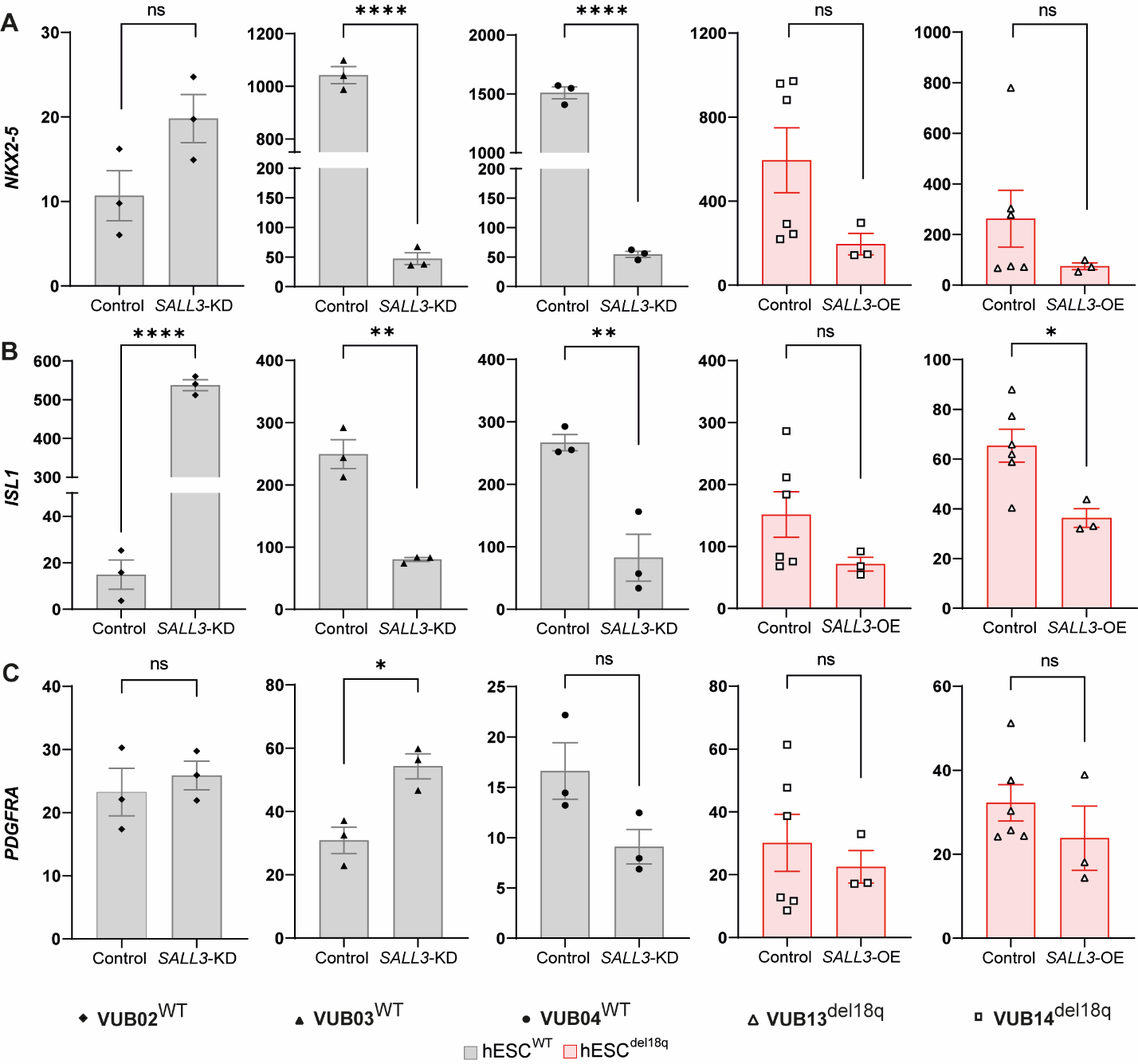

**Supplementary Figure 5. *SALL3* is not affecting the cells ability to generate mesoderm** (A-C) Relative mRNA expression for mesoderm markers measured by qRT-PCR (n = 3). Data are shown as the means ± SEM. Different patterns indicate different cell lines, and the horizontal bars with asterisks *, **, *** and **** represent statistical significance between samples at 5%, 1%, 0.1% and 0.01% respectively (unpaired t-test). *NKX2-5*(A), *ISL1*(B), *PDGFRA*(C).

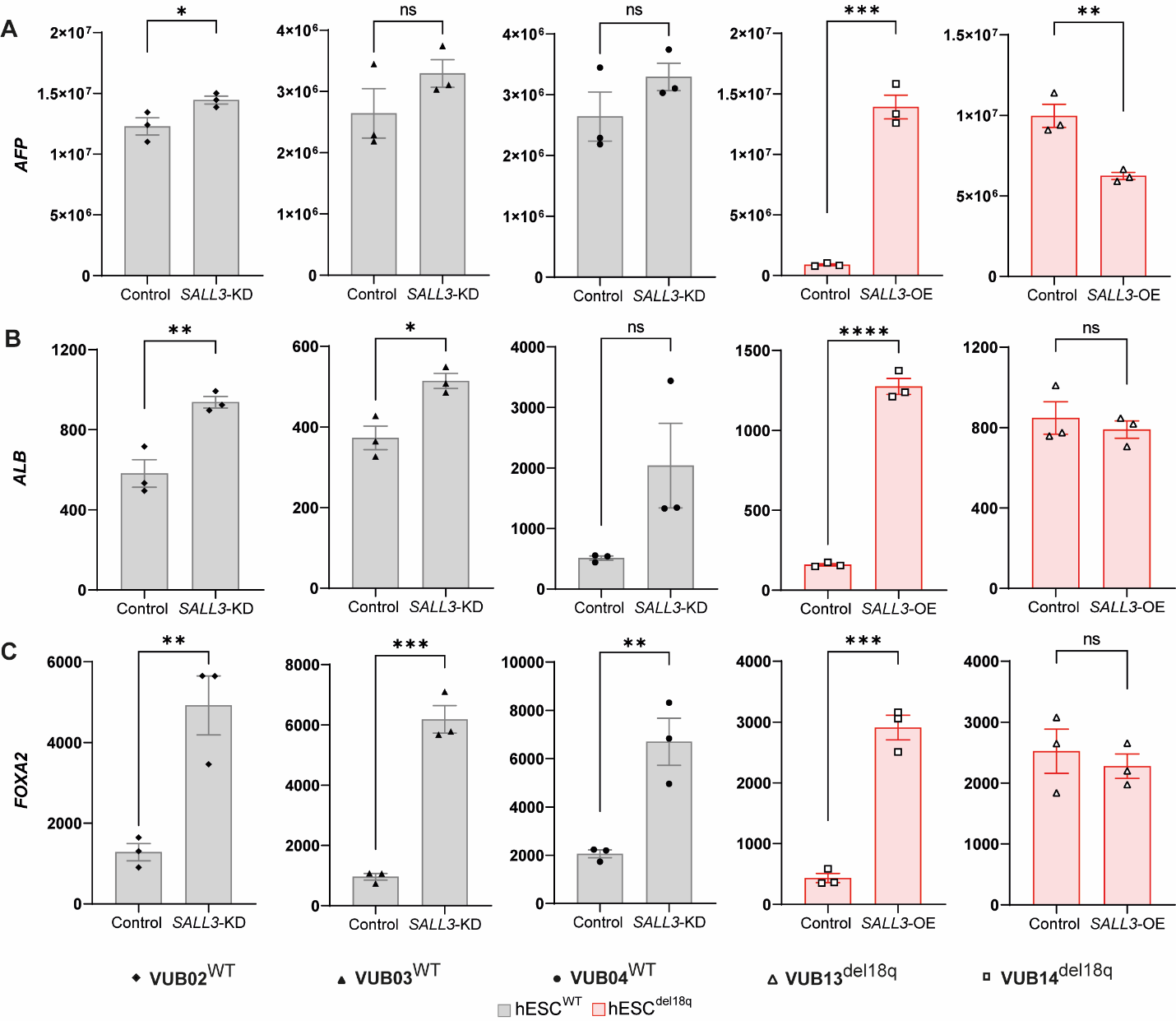

**Supplementary Figure 6. *SALL3* is affecting the cells ability to generate endoderm** (A-C) Relative mRNA expression for endoderm markers (n = 3) by qRT-PCR. Data are shown as the means ± SEM. Different patterns indicate different cell lines, and the horizontal bars with asterisks *, **, *** and **** represent statistical significance between samples at 5%, 1%, 0.1% and 0.01% respectively (unpaired t-test). *AFP*(A), *ALB*(B), *FOXA2*(C).

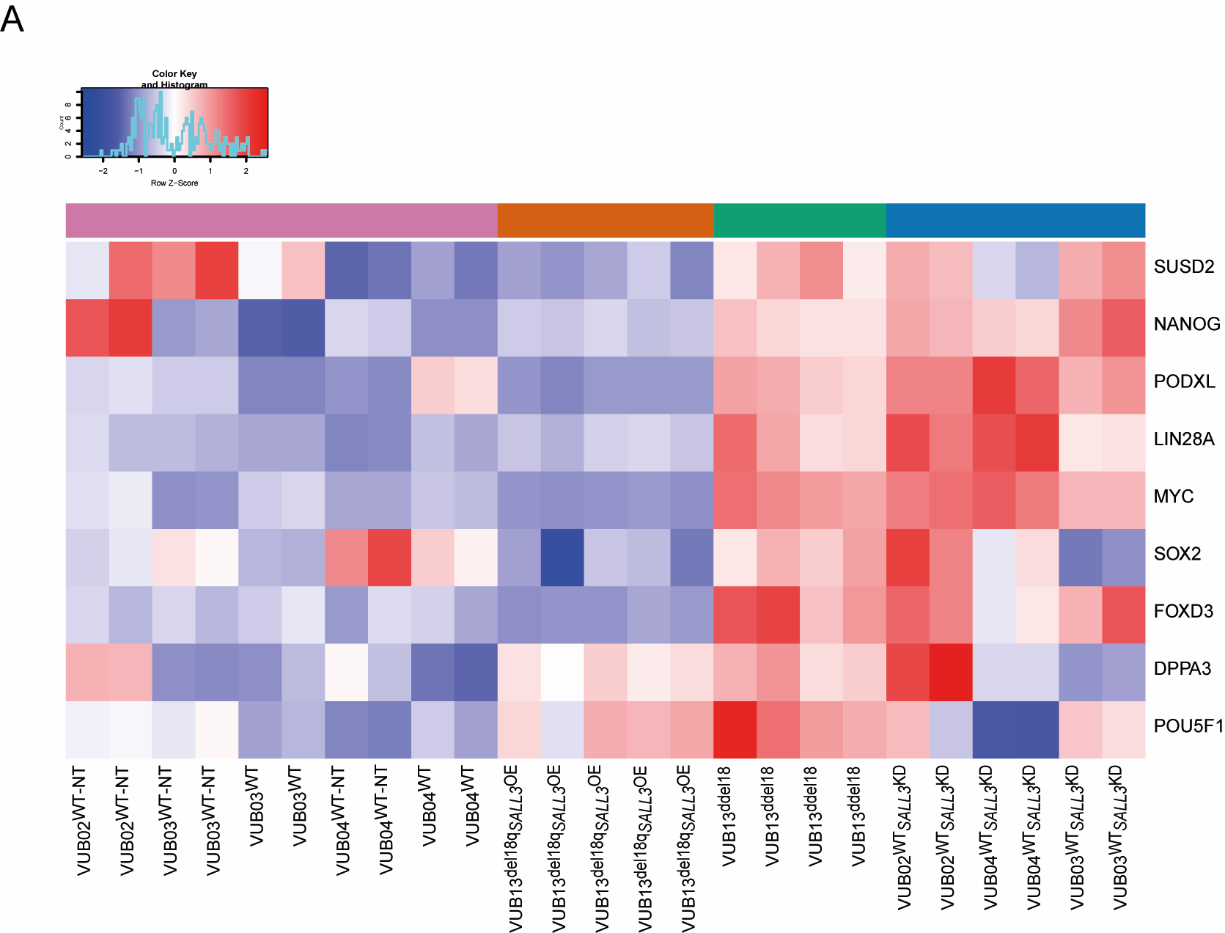

**Supplementary Figure 7. Heatmap of pluripotency-associated genes**

Supplementary tables 2, 3 and 4 are provided as excel files:

**Supplementary Table 2. Core set of 191 genes that are the most strongly regulated by *SALL3* in hESCs, both by the loss of a copy of the gene itself and by the modulation of its expression**

**Supplementary Table 3. Pathways enriched in the C2 and H libraries**

**Supplementary Table 4. Overlap in pathways enriched in VUB13^del18q^, hESC^WT_^*^SALL3^*^KD^ and VUB13^del18q_^*^SALL3^*^OE^**

**Supplementary table 5**

**TaqMan Gene Expression Assays**

| **Gene** | **Supplier** | **Catalog number** |
| --- | --- | --- |
| *GUSB* | Applied Biosystems™ | 4326320E |
| *PAX6* | Thermo Scientific | Hs0024087 |
| *SOX1* | Thermo Scientific | Hs01057642 |
| *NES* | Thermo Scientific | Hs04187831 |
| *AFP* | Thermo Scientific | Hs00173490 |
| *HNF4A* | Thermo Scientific | Hs00604435 |
| *FOXA2* | Thermo Scientific | Hs00232764 |
| *ALB* | Thermo Scientific | Hs00609411 |
| *GATA4* | Thermo Scientific | Hs01034629 |
| *NKX2-5* | Thermo Scientific | Hs00231763 |
| *PDGFRA* | Thermo Scientific | Hs00998018 |
| *ISL1* | Thermo Scientific | Hs00158126 |
| *POU5F1* | Thermo Scientific | Hs00742896 |
| *NANOG* | Thermo Scientific | Hs02387400 |

**TaqMan Copy Number Variation Assay**

| **Gene** | **Supplier** | **Catalog number** | **Chromosome** |
| --- | --- | --- | --- |
| RNaseP | Thermo Scientific | 4403326 | reference |
| ID1 | Thermo Scientific | Hs01892845 | 20q |
| KIF14 | Thermo Scientific | Hs00637799 | 1q |
| NANOG | Thermo Scientific | Hs03820140 | 12p |
| NMT1 | Thermo Scientific | Hs0550678 | 17q |

**Supplementary table 6**

**Primary Antibodies**

| **Gene** | **Species** | **Supplier** | **Catalog number** | **Dilution** |
| --- | --- | --- | --- | --- |
| PAX6 | Mouse | Abcam | ab78545 | 1:200 |
| HNF4A | Mouse | Santa Cruz Biotechnology | Sc-374229 | 1:200 |
| GATA4 | Rabbit | Cell Signaling Technology | 36966S | 1:200 |
| POU5F1 | Rabbit | Cell Signaling Technology | 2840S | 1:400 |
| POU5F1 | Mouse | Santa Cruz Biotechnology | sc-5279 | 1:200 |

**Secondary Antibodies**

| **Gene** | **Conjugate** | **Supplier** | **Catalog number** | **Dilution** |
| --- | --- | --- | --- | --- |
| Donkey anti Mouse | Alexa Fluor 488 | Thermo Fischer Scientific | A-21202 | 1:200 |
| Donkey anti Mouse | Alexa Fluor 594 | Thermo Fischer Scientific | A-21203 | 1:200 |
| Donkey anti Rabbit | Alexa Fluor 488 | Thermo Fischer Scientific | A-21206 | 1:200 |
| Donkey anti Rabbit | Alexa Fluor 594 | Thermo Fischer Scientific | A-21207 | 1:200 |
| Hoechst 33342 |  | Invitrogen | H3570 | 1:2000 |
